# Developmental Hypoxia Increases Susceptibility to Cardiac Ventricular Arrhythmias in Adult Offspring

**DOI:** 10.64898/2026.01.22.701057

**Authors:** Mitchell C Lock, Kerri LM Smith, Aga Swiderska, Hayat Baba, Andrew Silverwood, Julia Dyba, Olga V Patey, Youguo Niu, Sage G Ford, Freja Steinke, Katherine Dibb, Andrew W Trafford, Dino A Giussani, Gina LJ Galli

## Abstract

Ventricular arrhythmias are the leading cause of sudden cardiac death. It is well-established that environmental factors contribute to the origin and penetrance of ventricular arrhythmic disorders. However, to our knowledge, no studies have considered the role of the *intrauterine* environment. In this study, we investigated the long-term effects of fetal hypoxia on ventricular arrhythmia susceptibility. Pregnant Wistar rats were assigned to normoxia (21% O_2_) or hypoxia (13% O_2_ between gestational days 6-20), and offspring were raised to 6 months. Hearts were isolated and loaded with the fluorescent calcium- and voltage-sensitive indicators, Rhod-2 and RH237, respectively. Optical mapping was performed while the left ventricle was burst paced (10-20hz) to induce arrhythmias. Hearts isolated from adult offspring exposed to fetal hypoxia were more susceptible to arrythmia during burst pacing, compared to controls. This phenotype was associated with prolonged Ca^2+^ transients and action potentials, an increased frequency of Ca^2+^ waves and delayed after depolarisations, as well as lower gene and protein expression of the sarcoplasmic reticulum Ca^2+^ ATPase. Collectively, our data shows that fetal hypoxia can programme ventricular arrhythmia sensitivity in adulthood, driven by abnormalities in excitation-contraction coupling. This is the first evidence that some ventricular arrhythmias may have a developmental origin, highlighting pregnancy as a potential window for early preventive intervention.

## INTRODUCTION

Sustained ventricular arrhythmias are the leading cause of sudden cardiac death in the UK and Western world^1^. Treatment of these conditions is challenging because the origin of ventricular arrhythmias is often unrecognised until the person dies. Most ventricular arrhythmias occur in patients with structural heart diseases, such as ischemic and dilated cardiomyopathies^2^. However, some arrhythmias are found in individuals without any discernible cardiac abnormalities^3^. Identifying the risk factors that promote arrhythmia is crucial to implement interventional strategies that prevent arrhythmic death.

Genetics, lifestyle choices and environmental factors undoubtedly contribute to the origin^4–6^ and penetrance^7^ of ventricular arrhythmic disorders. However, to our knowledge, no studies have considered the role of the *intrauterine* environment. The seminal work from Barker in the 1980’s demonstrated that adverse events during pregnancy are strongly correlated with heart disease in offspring during adulthood. Since then, numerous maternal factors and pathologies have been linked with offspring cardiovascular disease^8–20^. Particularly damaging are conditions that reduce fetal oxygen supply (fetal hypoxia), such as preeclampsia, placental insufficiency, anaemia and high altitude pregnancies. Children born from these pregnancies have a range of cardiovascular abnormalities, including higher systolic and diastolic blood pressure, increased relative cardiac wall thickness, reduced left ventricular end diastolic volume and left ventricular diastolic dysfunction^8,9,14,21,22^. They are also more likely to develop early-onset cardiovascular disease, with two studies confirming an increased risk of arrhythmia in individuals from preeclamptic pregnancies^11,16^. Nevertheless, the link between fetal hypoxia and offspring arrhythmia sensitivity has never been explored.

Importantly, controlled studies from our laboratories^23–27^ and others (reviewed in ^28,29^) have shown that animal models of fetal hypoxia can recapitulate the cardiac phenotype from preeclamptic offspring, suggesting that fetal oxygen deprivation is the major factor driving cardiovascular dysfunction. Indeed, adult rodents and sheep from hypoxic pregnancies are hypertensive and possess cardiac abnormalities including ventricular hypertrophy, diastolic dysfunction and sympathetic dominance^30–34^. More recently, we have shown these abnormalities are partly mediated by dysfunctional ventricular myocyte Ca^2+^ handling, which is a well-established mechanism of ventricular arrhythmogenesis^35^. Therefore, it is possible that abnormal Ca²⁺ handling is the mechanistic link between preeclampsia and elevated risk of offspring arrhythmia in adulthood.

Several mechanisms can lead to Ca^2+^ mismanagement at the cellular level that drive ventricular arrhythmias (reviewed in^36^). Under normal conditions, an action potential (AP) generated from the sino atrial node propagates through the heart and activates L-type Ca^2+^ channels on the ventricular myocyte sarcolemmal membrane (for a review, see). This leads to Ca^2+^ entry which triggers a larger Ca^2+^ release from the ryanodine receptor (RyR) on the sarcoplasmic reticulum (SR) in a process known as Ca^2+^-induced Ca^2+^ release (CICR). Together, these pathways raise cytosolic free Ca^2+^ ([Ca^2+^]_i_) and promote Ca^2+^ binding to the myofilaments causing myocyte contraction. For relaxation, [Ca^2+^]_i_ is removed from the cytosol by the sarcoplasmic reticulum Ca^2+^-ATPase (SERCA2A) and the sarcolemmal Na^+^/Ca^2+^ exchanger (NCX). Reentry arrhythmias occur when AP propagation fails to terminate and tissue regions that have recovered are re-excited^37^. This mechanism commonly occurs in structural heart diseases where the AP travels around an obstacle, such as a fibrotic scar, or through cellular heterogeneities in the AP refractory period (the time taken for the AP to recover from excitation)^38^. Reentry mechanisms can give rise to both monomorphic ventricular tachycardia (MVT) and polymorphic ventricular tachycardia (PVT), which either terminate spontaneously or deteriorate to ventricular fibrillation (VF), causing cardiac arrest. In contrast to reentry, abnormal impulse formation, otherwise known as focal activity, mainly occurs with abnormal automaticity (e.g. altered pacemaker activity) or by the premature activation of cardiac tissue. The latter condition, known as triggered activity, is the result of early or delayed afterdepolarisations (EADs and DADs) which are abnormal depolarisations that interrupt the AP and cause premature ventricular contractions. EADs occur before terminal AP repolarization (phase 2 and phase 3) and they usually involve a prolonged AP duration. In contrast, DADs occur after the repolarization phase when membrane potential reaches resting levels. They originate from intracellular Ca^2+^ overload and the spontaneous release of Ca^2+^ from the SR, leading to propagating Ca^2+^ waves. Triggered activity usually gives rise to PVT which can degenerate into VT.

Our recent work has shown offspring from hypoxic pregnancies have smaller Ca^2+^ transients with a delayed diastolic phase^39^. When cytosolic Ca²⁺ remains elevated during diastole, it can promote DADs and EADs by increasing the likelihood of spontaneous calcium release from the SR, and prolonging the ventricular action potential duration, respectively. This is especially true when the ventricle is placed under physiological stress, such as high frequency pacing in the presence of adrenergic stimulation. In this study, we investigated the effects of fetal hypoxia on offspring arrhythmia sensitivity in a well-established rat model. We show that fetal hypoxia increases ventricular arrhythmia sensitivity in adulthood, which is associated with prolonged Ca^2+^ transients and action potentials, an increased frequency of Ca^2+^ waves and DADs, and lower gene and protein expression of SERCA-2A. To our knowledge, this is the first evidence that ventricular arrhythmia sensitivity can be programmed by the intrauterine environment.

## MATERIALS AND METHODS

### Animals, breeding and housing

This study was approved by the University of Manchester Ethical Review Board. All procedures were carried out in Manchester under the UK Animals (Scientific Procedures) 1986 Act. Wistar rats (Charles River Limited, UK) were delivered to the University of Manchester Biological Services Facility and housed in individually ventilated cages (IVC units, 21% O_2_, 70-80 air changes per hour) in rooms with controlled humidity (60%), controlled temperature (21 °C) and a 12:12 hour light-dark cycle with free access to food and water (Envigo). After one week of acclimatization, virgin female Wistar rats between 225-250g were paired individually with male Wistar rats. Cages were checked daily and the appearance of the copulatory plug was taken to be day 0 of pregnancy (term ∼22 days). On gestational day (GD) 6, pregnant rats were randomly allocated to two treatment groups: normoxia or hypoxia with n=8 per treatment group. Maternal weight, and food and water consumption were monitored throughout the experiment.

Pregnant rats assigned to the hypoxia group were placed inside a transparent hypoxic chamber (Coy Laboratory Products, Inc. USA), as previously described. The chamber consists of a clear PVC isolator attached to nitrogen and oxygen cylinders to precisely control the percentage of oxygen within the chamber. Carbon dioxide and atmospheric waste were scavenged by circulating the air through soda lime pellets and activated charcoal pellets, respectively (Sigma-Aldrich, USA). The chamber contained a hygrometer and thermometer for continuous monitoring of humidity and temperature whilst oxygen was monitored with the inbuilt calibrated oxygen sensor (Coy Laboratory Products, Inc. USA). Pregnancies undergoing maternal hypoxia were maintained at an inspired fraction of oxygen of 13% (± 0.1%) from GD6-GD20 and food and water consumption were recorded every 48 hrs.

At GD20, rats were returned to normal IVC cages for birth at GD23. At birth each litter was reduced to 8 pups (4 males and 4 females) to standardize nutritional intake. At weaning males and females were separated and housed as groups of 4 siblings until 16-19 weeks of age. Animals were humanely killed by CO_2_ inhalation followed by cervical spinal transection. At this time, animals from each litter were randomly assigned to undergo either; tissue collection, cardiac myocyte isolation, or optical mapping experiments.

### Tissue collection and cardiac myocyte isolation in adult offspring (4 months of age)

The body and major organs were weighed and tissue samples were collected from the left ventricle free wall. Samples were frozen in liquid nitrogen and stored at −80 °C for later analysis. Animals assigned to the cardiac myocyte isolation group were humanely killed with CO_2_ inhalation followed by cervical spinal transection. The heart was then quickly removed and cannulated on a langendorff apparatus retrogradely perfusing at ∼10 mL/min through the aorta with a nominally Ca^2+^-free solution for 5 minutes at 37°C (Ca^2+^-free solution [mmol/L] NaCl 134, Glucose 11.1, HEPES 10, KCl 4, MgSO_4_ 1.2, and NaH_2_PO_4_ 1.2, pH 7.34 with NaOH). Liberase TM (Roche, Switzerland) at 6 Wünsch units was added to 50ml Ca^2+^-free solution and perfused for 10-13 minutes until the heart was visibly flaccid. The perfusate was then changed to a taurine containing solution for 10 minutes (taurine Solution [mmol/L] NaCl 115, Glucose 11.1, HEPES 10, KCl 4, MgSO_4_ 1.2, NaH_2_PO_4_ 1.2, and Taurine 50, pH 7.34 with NaOH). The heart was then dissected, left ventricle and right ventricles were separated and cut into small pieces in a taurine solution and gently triturated to liberate single cells. The isolated cells were stored at room temperature in a taurine-containing solution until use.

### Measurement of collagen content

The heart was excised, and 2–3 mm thick slices were cut in the central region using a matrix slicer to obtain a transverse section of the right and left ventricles. The transverse sections of the heart were immersed in neutral buffered 10% formalin overnight, then transferred to phosphate buffer solution (PBS) before being embedded in paraffin and sectioned into 5 µm slices. Sections were stained using a Picrosirius Red staining, which involved deparaffinizing the slides in xylene and rehydrating them through a graded ethanol series (100%, 95%, 70%) to PBS. The sections were then incubated in 0.1% Sirius Red dissolved in a saturated picric acid solution for 1 hour, followed by two washes in 0.5% acetic acid to remove excess stain. The stained sections were dehydrated in 100% ethanol (three washes), cleared in xylene, and mounted for imaging. Interstitial collagen content was quantified by analyzing ten randomly selected regions per ventricle per section, while perivascular collagen was assessed in five to seven fields per section. Imaging was performed at 20x magnification using a single-slide scanner microscope (BX63) equipped with polarized light, capturing both bright-field and polarized images using Olympus cellSens software. Image analysis was conducted using ImageJ software, where the collagen-positive area in the polarized images was quantified and divided by the total tissue area in the corresponding bright-field image. The results were expressed as the percentage of total collagen area.

### Measurement of intracellular Ca^2+^ ([Ca^2+^]_i_)

Cells were loaded with the Ca^2+^ sensitive fluorescent indicator Fura-2 AM at 1µmol/L for 5 minutes in a Tyrode solution containing (mmol/L) NaCl 140, HEPES 10, Glucose 10, Probenecid 2, KCl 4, MgCl_2_ 1, CaCl_2_ 1, pH 7.34 with NaOH. Loaded cells were then pipetted onto a Nikon Eclipse TE2000-U microscope with a Nikon Plan Flour 40X Oil immersion objective lens (Japan) fitted with a MPRE8 inline heated tip (Cell MicroControls, USA) with constant perfusion of tyrode solution at 37 °C. Individual cardiac myocytes were located and simultaneously excited at 340 nm and 380 nm and field stimulated at 1 Hz. [Ca^2+^]_i_ was recorded for 5 minutes until Ca^2+^ transients were stabilized. The cellular response to β-Adrenergic stimulation was induced by exposure to 100nmol L^-1^ Isoprenaline Sulphate (Stockport Pharmaceuticals, Stepping Hill Hospital, UK). Ca^2+^ transients were analysed using Clampfit 11 software (Molecular Devices, USA).

### Calcium sparks

Isolated ventricular myocytes were loaded with the Ca²⁺ sensitive dye Fluo-5F AM (Invitrogen) for 10 minutes at RT, before resuspension in Tyrode solution (as above). Cells were allowed to de-esterify for 30 minutes at RT before experiments were performed. Calcium spark frequency was assessed using a Zeiss LSM7Live scanning confocal microscope (Zeiss, Germany), at 63x magnification. Zen software (Zeiss, Germany) was used for cell line scanning, with a line drawn parallel to the longitudinal axis of the cell, including the boundary of each cell edge. Cells were stimulated at 0.5 Hz, and visualised at a scan speed of 1000 lines per second. Cells were excited at 488 nm, and emission collected using a 525/50 band pass dichroic filter. Spark frequency was assessed under normal conditions (perfusion with Tyrode solution at 37°C) and in response to β-adrenergic stimulation (perfusion with isoprenaline sulphate [100 nM]). Analysis of calcium spark data was performed using MATLAB software, utilising an in-house code written by Dr Edward Hayter (Sparkinator). In brief, a cell profile was created by averaging pixels across time and smoothing after which the edges of the cell were automatically identified and cropped. The baseline was removed from each pixel and stimulated peaks were excluded by removing the first 800 ms following stimulation. Peaks in the remaining 1200 ms window were detected following application of a threshold, resulting in an overall normalised spark count (sparks/μm/sec). Spark counts were calculated over a five stimulation (or 10 second) window.

### Optical mapping experiments in adult offspring

At 4-5 months of age, animals were humanely killed and the heart was removed. The aorta was cannulated and quickly perfused with 5 ml ice cold oxygenated KH solution ([mmol/L] NaCl 119, NaHCO_3_ 25, KCl 4, KH_2_PO_4_ 1.2, MgCl_2_ 1, CaCl_2_.2H_2_O 1.8, D-glucose 10, buffered using 95% O_2_, 5% CO_2_) containing 1000U Heparin. The heart was then transferred onto a langendorff apparatus retrogradely perfusing at ∼5mL/min through the aorta at 37 °C and allowed to stabilise (heart rate >250 bpm) for 10 minutes. 200 µl blebbistatin solution (1mg/ml) was slowly injected into the perfusate via the drug port to inhibit myosin and hence contraction of the heart. 50 μl of Pluronic (20% Solution in DMSO) was then added to the perfusate and allowed to circulate for 10 mins. 100 μ1 Rhod-2AM (1 mg/ml) was slowly injected through the drug port over 5-10 mins, followed by 100 μl of RH237 (1.2 mg/ml). The heart was then paced using a MappingLab stimulation electrode gently inserted into the epicardial wall of the apex. ECG’s were recorded with electrodes and Ca^2+^ (Excitation/Emission: 540/585) and voltage (Excitation/Emission: 540:650) were imaged with a MappingLab OMS-PCIE-2002 Optical Mapping system using OMapRecord 5 software.

A custom burst pacing stimulation protocol was utilised to generate ventricular arrhythmias, leaving a three minute unstimulated resting phase in-between each stage to allow the heart to recover. Hearts were left to stabilise in sinus rhythm for 10 minutes, and then subjected to increasing burst pacing (5, 10, 12, 14, 16, 18, and 20 Hz) for three seconds, followed by sinus rhythm recording between each frequency. Isoprenaline was then added (100nmol L^-1^), and the burst pacing protocol was repeated. Arrhythmia was defined as sustained loss of 1:1 AV conduction with sustained arrhythmia for >1 minute monitored by pseudo-ECG. The percent change in diastolic calcium was calculated from a 1cm^2^ area of the left ventricle using Rhod2 fluorescence recordings (percentage change in Rhod2 fluorescence from sinus rhythm to the beginning of arrhythmia).

### Real-time PCR for target genes

Male offspring from Normoxic (n=8) and Hypoxic (n=8) pregnancy were used for qRT-PCR experiments. All essential information regarding the qRT-PCR procedure is included as per the MIQE guidelines. Total RNA was extracted from frozen left ventricle tissue for each animal using QIAzol Lysis Reagent solution and QIAgen RNeasy purification columns, as per manufacturer guidelines (Qiagen, Germany). Total RNA was quantified by spectrophotometric measurements at 260 and 280 nm in a NanoDrop Lite Spectrophotometer (Thermo Fisher Scientific). The 260/280 nm ratio results were less than 2.1 and greater than 1.9 and therefore acceptable for qRT-PCR. cDNA was synthesised using Superscript IV First Strand Synthesis System (Invitrogen, USA) using 1 µg of total RNA, random hexamers, dNTP, DTT and Superscript IV in a final volume of 20 µL, as per the manufacturer’s guidelines in a C1000 thermocycler (Biorad, USA). Controls containing either no RNA transcript or no Superscript IV were used to test for reagent contamination and genomic DNA contamination, respectively. The geNorm component of qbaseplus 2.0 software (Biogazelle, Belgium) was used to determine the most stable reference genes from a panel of candidate reference genes and the minimum number of reference genes required to calculate a stable normalisation factor. For qRT-PCR data output normalisation, two stable reference genes RPL4 and RPL32 (Table 1) were run in parallel with all target genes, as previously described ^40^. A selection of genes was chosen *a priori* to investigate components of the excitation-contraction coupling pathway. Primers were validated and optimised as previously described ^41–43^. Relative expression of target genes (Table 1) were measured by Power SYBR™ Green PCR Master Mix (Thermo Fisher Scientific) in a final volume of 9 µL on a ViiA7 Fast Real-time PCR system (Applied Biosystems, USA). Each qRT-PCR well contained 6 µL SYBR Green Master Mix, 1 µL each of forward and reverse primer mixed with H_2_O to obtain final primer concentrations and 1 µL of diluted cDNA. Each sample was run in triplicate for target and reference genes. The abundance of each transcript relative to the abundance of stable reference genes ^44^ was calculated using DataAssist 3.0 analysis software (Applied Biosystems, USA) and expressed as mRNA mean normalised expression (MNE) ± SD. Outliers were determined using the ROUT method and removed from the analysis (GraphPad Prism 8, USA).

**Table 1:**
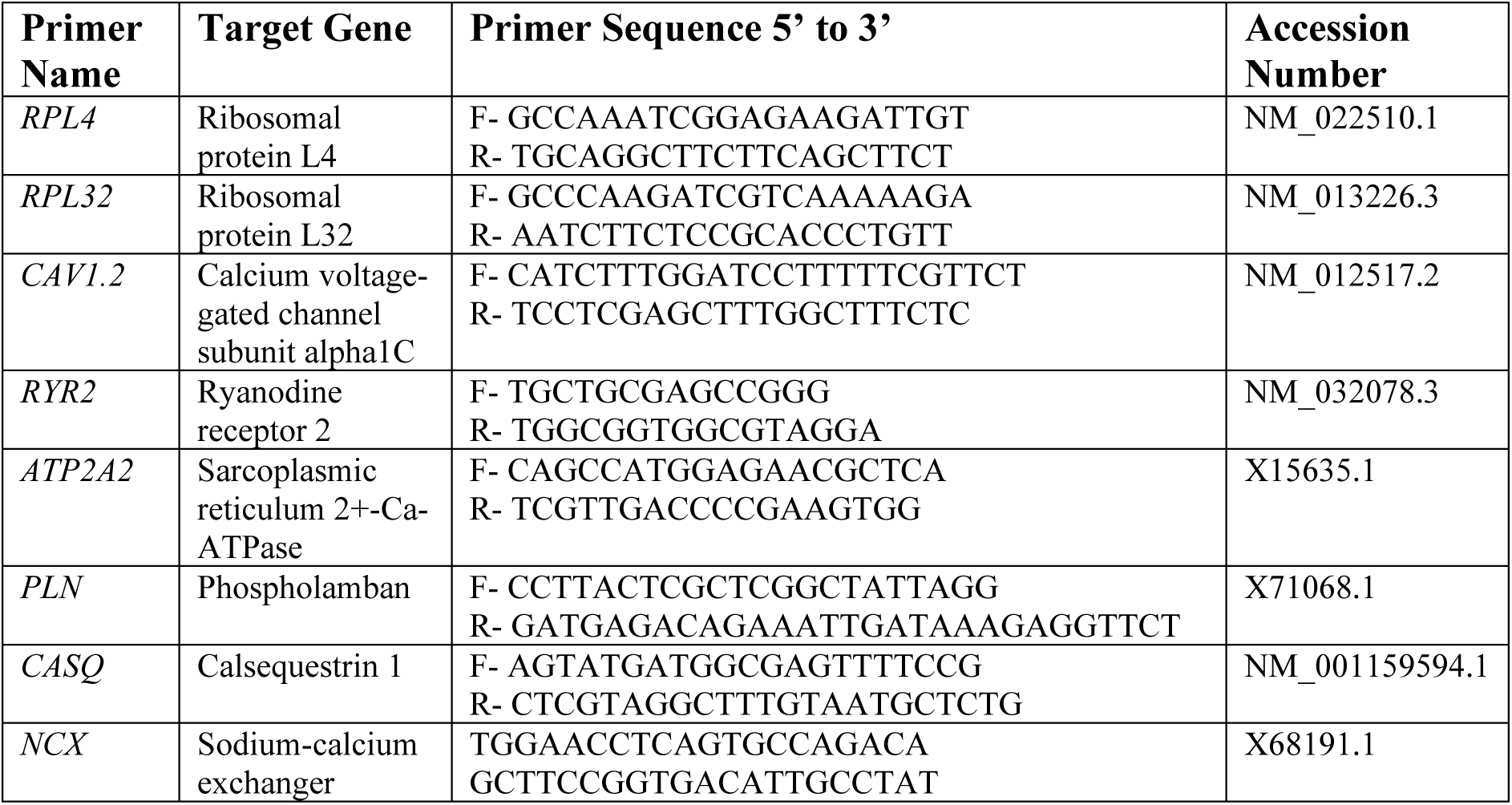
qRT-PCR Primer Information.

**Table 2:** Isolated cell and optical mapping measures.

| Experiment | Measure | Males |  | Females |  | P Value<br>H=Hypoxia<br>S=Sex<br>I=Interaction |
| --- | --- | --- | --- | --- | --- | --- |
|  |  | Normoxia | Hypoxia | Normoxia | Hypoxia |  |
| Isolated cells after isoprenaline treatment | Spontaneous calcium release events (per minute) | 19.16±19.44 (n=28) | 43.82±8.92 (n=15) | 14.81±14.38 (n=27) | 35.81±12.63 (n=15) | H:<**0.0001** S:0.0786 I:0.5994 |
|  | Amplitude of calcium waves (ratio 340/380) | 0.0407±0.0125 (n=14) | 0.0520±0.0164 (n=19) | 0.0386±0.0118 (n=15) | 0.0460±0.0140 (n=19) | H:**0.0092** S:0.2447 I:0.5756 |
| Sinus rhythm optical mapping | DT20 (ms) | 21.51±6.26 (n=7) | 24.40±2.82 (n=8) | 21.07±7.35 (n=7) | 25.74±9.48 (n=8) | H:0.1477 S:0.8587 I:0.7284 |
|  | DT50 (ms) | 45.054±12.65 (n=7) | 58.46±5.44 (n=8) | 47.47±17.22 (n=7) | 55.14±16.60 (n=8) | H:**0.0453** S:0.9290 I:0.5724 |
|  | DT90 (ms) | 82.44±20.42 (n=7) | 106.46±10.13 (n=8) | 86.60±31.13 (n=7) | 101.67±29.34 (n=8) | H:**0.0354** S:0.9720 I:0.6157 |
|  | Decay time constant (Tau) | 24.05±11.72 (n=7) | 35.03±12.61 (n=8) | 21.75±11.89 (n=6) | 25.25±11.32 (n=7) | H:0.1230 S:0.1956 I:0.4174 |
|  | Time to peak (ms) | 4.42±1.68 (n=7) | 4.17±1.12 (n=8) | 4.95±0.69 (n=7) | 4.73±0.98 (n=8) | H:0.5851 S:0.2109 I:0.9767 |
|  | APD20 (ms) | 8.23±0.95 (n=7) | 9.08±2.99 (n=8) | 8.93±2.28 (n=8) | 10.22±2.39 (n=8) | H:0.2109 S:0.2813 I:0.7964 |
|  | APD50 (ms) | 17.79±2.87 (n=7) | 21.00±6.26 (n=8) | 19.65±4.31 (n=8) | 23.73±5.38 (n=8) | H:**0.0498** S:0.2075 I:0.8087 |
|  | APD90 (ms) | 44.09±9.76 (n=7) | 56.40±13.92 (n=8) | 44.24±7.55 (n=8) | 57.80±9.41 (n=8) | H:**0.0019** S:0.8390 I:0.8681 |
|  | CV (mm/ms) | 5.38±2.02 (n=7) | 4.52±2.23 (n=7) | 4.94±1.35 (n=8) | 3.21±0.97 (n=7) | H:0.0518 S:0.1783 I:0.4966 |
| 5Hz stimulated optical mapping | DT20 (ms) | 20.17±5.60 (n=7) | 24.37±4.72 (n=7) | 22.78±3.79 (n=7) | 21.55±5.03 (n=7) | H:0.4233 S:0.9556 I:0.1494 |
|  | DT50 (ms) | 41.66±11.04 (n=7) | 50.08±7.03 (n=7) | 47.80±5.54 (n=7) | 45.12±6.30 (n=7) | H:0.3384 S:0.8444 I:0.0711 |
|  | DT90 (ms) | 74.50±20.79 (n=7) | 91.33±4.99 (n=7) | 84.27±8.72 (n=7) | 83.86±9.80 (n=7) | H:0.0962 S:0.8103 I:0.0817 |
|  | Decay time constant (Tau) | 11.74±1.68 (n=7) | 16.84±3.95 (n=8) | 13.25±2.34 (n=7) | 12.96±3.61 (n=8) | H:**0.0435** S:0.3045 I:**0.0254** |
|  | Time to peak (ms) | 2.59±0.29 (n=7) | 2.57±0.19 (n=7) | 2.71±0.28 (n=7) | 2.60±0.30 (n=8) | H:0.5222 S:0.4598 I:0.6335 |
|  | APD20 (ms) | 10.06±1.70 (n=7) | 11.58±2.50 (n=7) | 10.58±2.48 (n=6) | 13.06±3.22 (n=6) | H:0.0543 S:0.3215 I:0.6295 |
|  | APD50 (ms) | 24.90±6.52 (n=7) | 26.85±7.59 (n=7) | 24.26±6.34 (n=6) | 29.75±9.43 (n=6) | H:0.2220 S:0.7073 I:0.5555 |
|  | APD90 (ms) | 50.60±11.20 (n=7) | 65.29±11.53 (n=7) | 53.49±14.14 (n=6) | 59.70±15.21 (n=6) | H:0.0528 S:0.7941 I:0.4155 |
|  | CV (mm/ms) | 2.10±0.96 (n=7) | 1.47±0.53 (n=7) | 1.60±0.37 (n=8) | 1.71±0.58 (n=8) | H:0.2745 S:0.5932 I:0.1264 |
| 10Hz burst pacing optical mapping | DT20 (ms) | 10.50±1.93 (n=7) | 11.16±1.80 (n=7) | 11.29±1.34 (n=8) | 11.73±1.90 (n=8) | H:**0.0019** S:0.7335 I:0.8057 |
|  | DT50 (ms) | 21.00±3.02 (n=7) | 28.07±2.68 (n=7) | 23.76±3.85 (n=8) | 24.00±3.67 (n=8) | H:**0.0031** S:0.6821 I:0.7685 |
|  | DT90 (ms) | 41.11±6.78 (n=7) | 51.68±3.56 (n=7) | 43.57±6.32 (n=8) | 43.44±6.04 (n=8) | H:**0.0118** S:0.7021 I:0.9759 |
|  | Decay time constant (Tau) | 4.24±1.27 (n=7) | 5.79±2.43 (n=8) | 4.09±0.78 (n=8) | 4.95±0.93 (n=8) | H:**0.0349** S:0.3712 I:0.5369 |
|  | Time to peak (ms) | 2.44±1.29 (n=7) | 3.14±1.19 (n=8) | 3.67±1.43 (n=7) | 2.96±1.38 (n=8) | H:0.9920 S:0.2847 I:0.1571 |
|  | APD20 (ms) | 10.50±1.93 (n=7) | 11.16±1.80 (n=7) | 11.29±1.34 (n=8) | 11.73±1.90 (n=8) | H:0.4000 S:0.2968 I:0.8616 |
|  | APD50 (ms) | 21.00±3.02 (n=7) | 28.07±2.68 (n=7) | 23.76±3.85 (n=8) | 24.00±3.67 (n=8) | H:**0.0065** S:0.5986 I:**0.0103** |
|  | APD90 (ms) | 41.11±6.77 (n=7) | 51.68±3.56 (n=7) | 43.57±6.32 (n=8) | 43.44±6.04 (n=8) | H:**0.0217** S:0.1883 I:**0.0189** |
|  | CV (mm/ms) | 1.56±0.59 (n=7) | 1.56±0.56 (n=7) | 1.68±0.65 (n=8) | 1.32±0.27 (n=8) | H:0.3632 S:0.7488 I:0.3784 |

### Western Blot

Protein was extracted from frozen left ventricle tissue for each animal. 50mg tissue was resuspended in RIPA buffer containing 1% Protease and Phosphatase Inhibitor Cocktail (Sigma) to generate a 10% homogenate. Tissue was homogenised in a Precellys Evolution at 4°C, and homogenate centrifuged at 10000 x g for 10 minutes at 4°C. Supernatant was collected, and protein concentration measured using a DC protein assay (Bio-Rad). Supernatant was aliquoted to avoid freeze/thaw cycles and stored at −80°C until use. For RYR2 and phospho-RYR2-S2808, protein was diluted in RIPA buffer, reduced at RT for 15 minutes and loaded onto a 3-8% Tris-Acetate gel. For all other proteins, protein was diluted in RIPA buffer, reduced and denatured by boiling for 10 minutes, and loaded onto a 4-12% Bis-Tris gel. For SERCA2 and phospholamban (PLN), 2 µg of protein was loaded. For all other proteins, 10 µg of protein was loaded. Following electrophoresis, proteins were transferred to a nitrocellulose membrane using a Trans-Blot Turbo system (BioRad). Membranes were either blocked with 5% bovine serum albumin (BSA) in TBS-T (phospho-PLN-T17, phospho-RYR2-S2808 and phospho-cardiac Troponin I [cTnI]), or with 5% milk in TBS-T (PLN, cTnI, Calsequestrin-1, SERCA2, RYR2, NCX1 and LTCC). Membranes were incubated with primary antibodies (PLN [1/5000 dilution] – MA3-922, InVitrogen; phospho-PLN-T17 [1/1000 dilution] – AP0910, ABclonal; cTnI [1/1000 dilution] – MCA1208, Bio-Rad; phospho-cTnI [1/1000 dilution] – MCA2780, Bio-Rad; Calsequestrin-1 [1/5000 dilution] – PA1-913, InVitrogen; SERCA2 [1/2000 dilution] – ab3625, Abcam; NCX1 [1/1000 dilution] – R3F1, Swant; LTCC [1/1000 dilution] – ab2864, Abcam; RYR2 [1/1000 dilution] – ab302716, Abcam; phospho-RYR2-S2808 [1/1000 dilution] – PA5-105712, InVitrogen) overnight at 4°C, before secondary antibody incubation (IRDye® 800CW Goat anti-mouse for PLN, cTnI, phospho-cTnI, NCX1 and LTCC, IRDye® 800CW goat anti-rabbit for phospho-PLN-T17, Calsequestrin-1, SERCA2, RYR2 and phosphor-RYR2-S2808, both 1/20 000 dilution) at room temperature for one hour. Protein abundance was visualised by fluorescence (Licor-Odyssey CLx, ex/em 778/795 nm), and normalised to total lane protein (membrane stained using Revert 700 Total Protein Stain, Licor) and an internal control for inter-membrane normalisation. Each sample was repeated in triplicate across three individual gels, and results averaged. Blots were imaged and analysed using Image Studio (v5.2). Uncropped representative images of blots are shown in Supplementary Figure S3.

### Statistical Analysis

Statistical analyses were performed in Graphpad Prism 9 (Graphpad Software Inc., USA) or SPSS Statistics (IBM) for nested analyses. All analyses were assessed for normal distribution of data and a *P<0.05* was considered significant. Outliers from the qRT-PCR data were determined using the ROUT method (Q=1%) and removed from the analysis. Analysis of normoxia *vs.* hypoxia and male *vs.* female was performed using a 2-way ANOVA to test for an effect of hypoxia, sex and the interaction of these main effects. Since some isolated cells were from the same animal; linear mixed modelling was performed to account for the nested (clustered) design of the experiment. If a significant interaction was determined, the data were divided by treatment and assessed with a Tukey’s post-hoc test. As only male offspring were used for qRT-PCR, Western blot and calcium sparks experiments, these were analysed by a Student’s t-test (Graphpad).

## RESULTS

### Fetal hypoxia increases offspring cardiac arrhythmia susceptibility

Control rat hearts either remained rhythmic throughout the entire protocol (n=5) or only experienced arrhythmia at 20Hz (n=1), after isoprenaline infusion (n=2) or with a combination of isoprenaline and burst pacing (n=6) (Fig. 1 black bars; average stage of the protocol where arrhythmias started was ± 3.72). In contrast, the hearts of adult offspring from hypoxic pregnancies were significantly more sensitive to ventricular arrhythmia (P<0.001), both in sinus rhythm and under physiological stress (Fig. 1, red bars). Indeed, most males from hypoxic pregnancies developed arrhythmia in the absence of isoprenaline at 12-20Hz (n=4), or after isoprenaline infusion (n=1) (Fig 1, average stage of protocol = ± 3.72). Some males (n=2) even developed arrhythmias in sinus rhythm or at low physiological pacing frequencies (5-10Hz). Females from hypoxic pregnancies were also more susceptible to arrhythmia than controls (P=0.004, average stage ± 3.06 stages), but less than their male counterparts (P<0.0001), with arrhythmias only developing at 14-16Hz (n=3) or with a combination of isoprenaline and burst pacing (n=4). We observed examples of both MVT and PVT (Fig. 2A-C) during the burst pacing protocols, but MVTs always developed into PVT’s if the hearts did not recover after the burst protocol (Fig. 2D).

**Figure 1:**
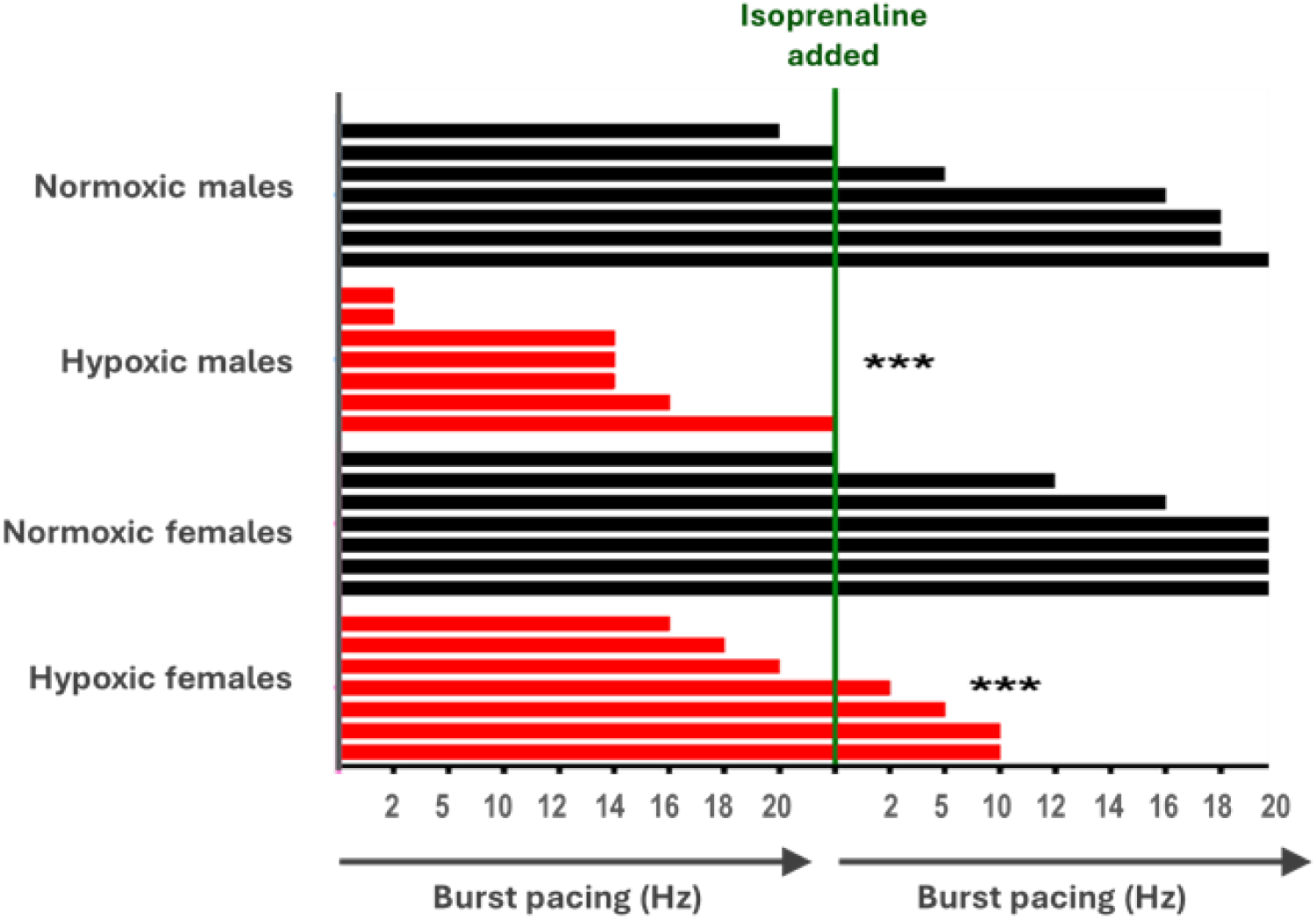
Fetal hypoxia increases sensitivity in 6-month rat offspring. Isolated rat hearts from normoxic pregnancies (black bars, n=7 per sex; 21% O_2_) and hypoxic pregnancies (red bars, n=7 per sex; 13% O_2_ from gestational days 6-20) were exposed to burst pacing (from 2 to 20 Hz), Isoprenaline was added and the burst pacing protocol was repeated. Each bar represents a heart from a single animal, the end of the bar signifies the stage of the pacing protocol that the heart became arrhythmic. Normoxic Males (n=7), Normoxia Females (n=7), Hypoxia Males (n=7), Hypoxia Females (n=7).

**Figure 2:**
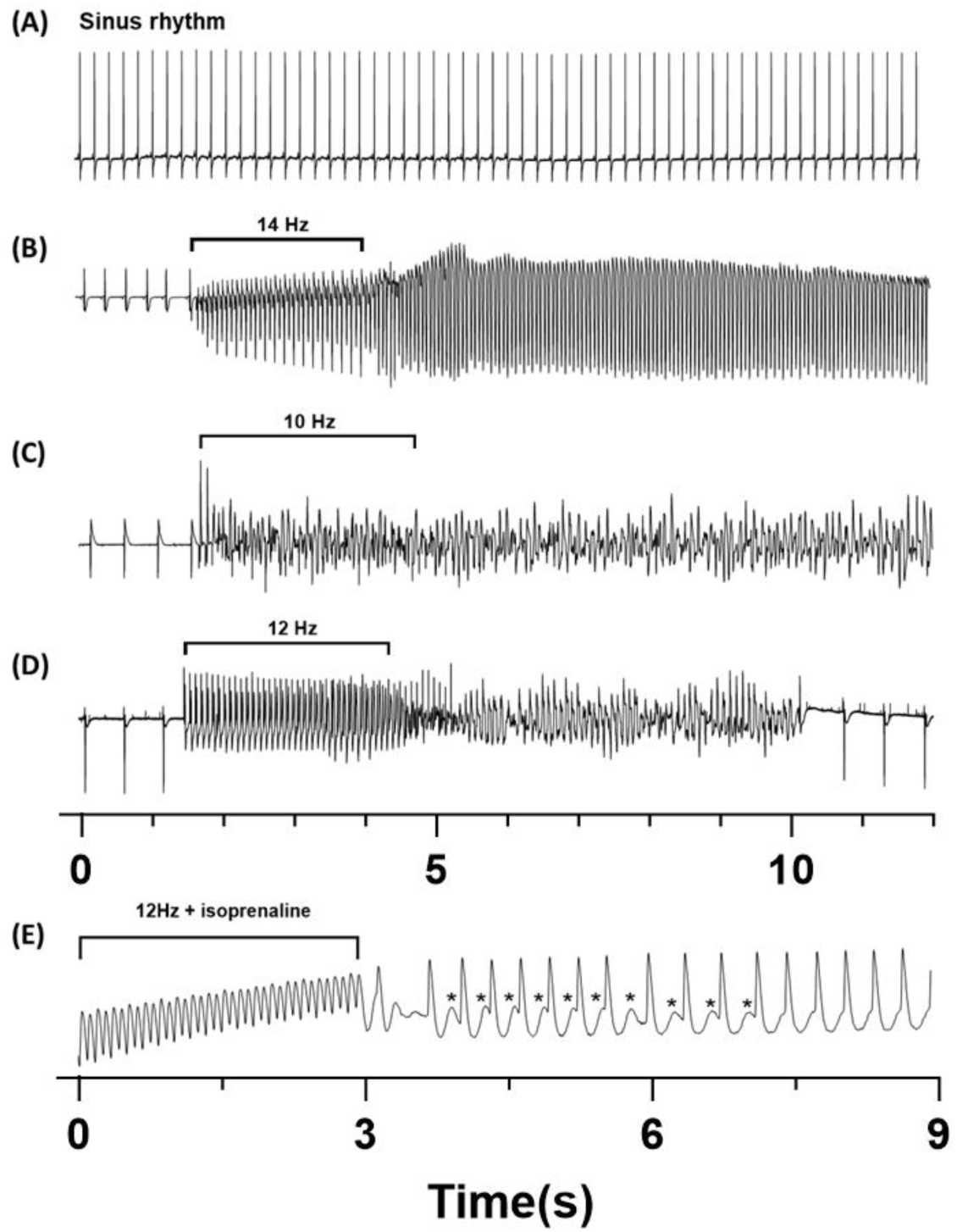
Original traces of ECG and calcium fluorescence from adult male offspring from hypoxic pregnancies. Optical mapping was used to measure intracellular calcium and electrocardiograms (ECG) in isolated rat hearts. The traces in this figure are all from adult male offspring from hypoxic pregnancies. ECG in sinus rhythm (A) and after a burst pacing protocol (B-D) where the ventricle is stimulated for 3 seconds at frequencies ranging between 5 to 20Hz, and then stimulation is switched off. In some circumstances, the ventricles went into monomorphic ventricular tachycardia after burst pacing (B), while others went into polymorphic ventricular tachycardia (C) or a combination of the two (D). In some cases, hearts eventually recovered from the protocol and returned to sinus rhythm (D). Panel (E) shows intracellular calcium in a heart that was burst paced at 12Hz in the presence of isoprenaline. An asterisk is placed next to calcium transient with spontaneous calcium release during diastole (calcium waves).

### Collagen content and conduction velocity

While there was a strong trend (P=0.0518) towards a prolonged conduction velocity with fetal hypoxia, there were no significant differences between experimental groups in this parameter (Fig. 3). Likewise, there was no difference in collagen content between experimental groups in the left or right ventricle (Fig. 4).

**Figure 3:**
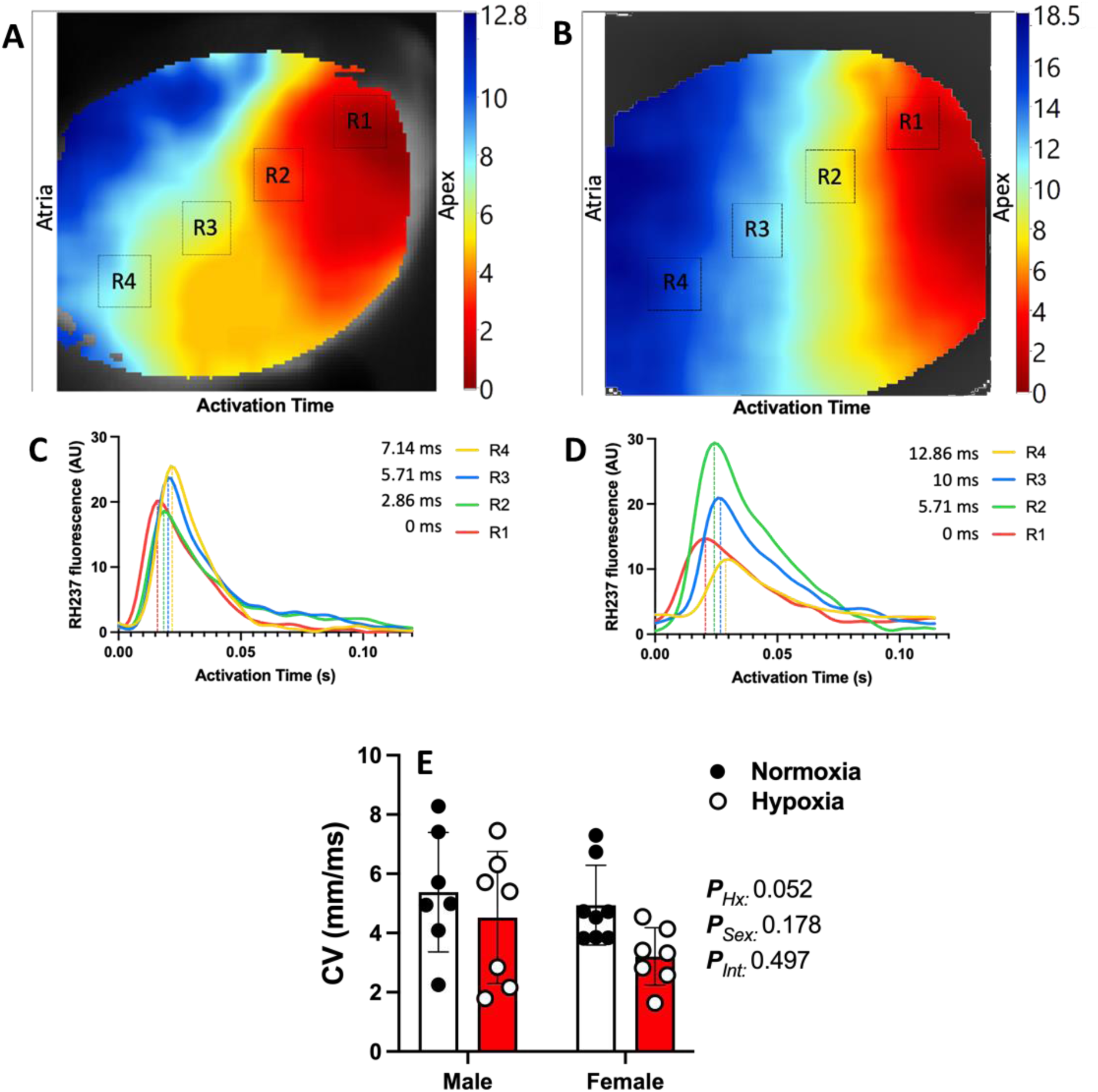
Conduction velocity measures across the left ventricle of hearts from offspring of normoxic or hypoxic pregnancy. (A, B) Conduction velocity representative heart images from the left ventricle of Control and Hypoxic offspring respectively. (C, D) Corresponding activation times for each region within the left ventricle: R1; Region 1, R2; Region 2, R3; Region 3, R4; Region 4. (E) Average conduction velocity across the left ventricle in Normoxic and Hypoxic offspring of hypoxic pregnancies.

**Figure 4:**
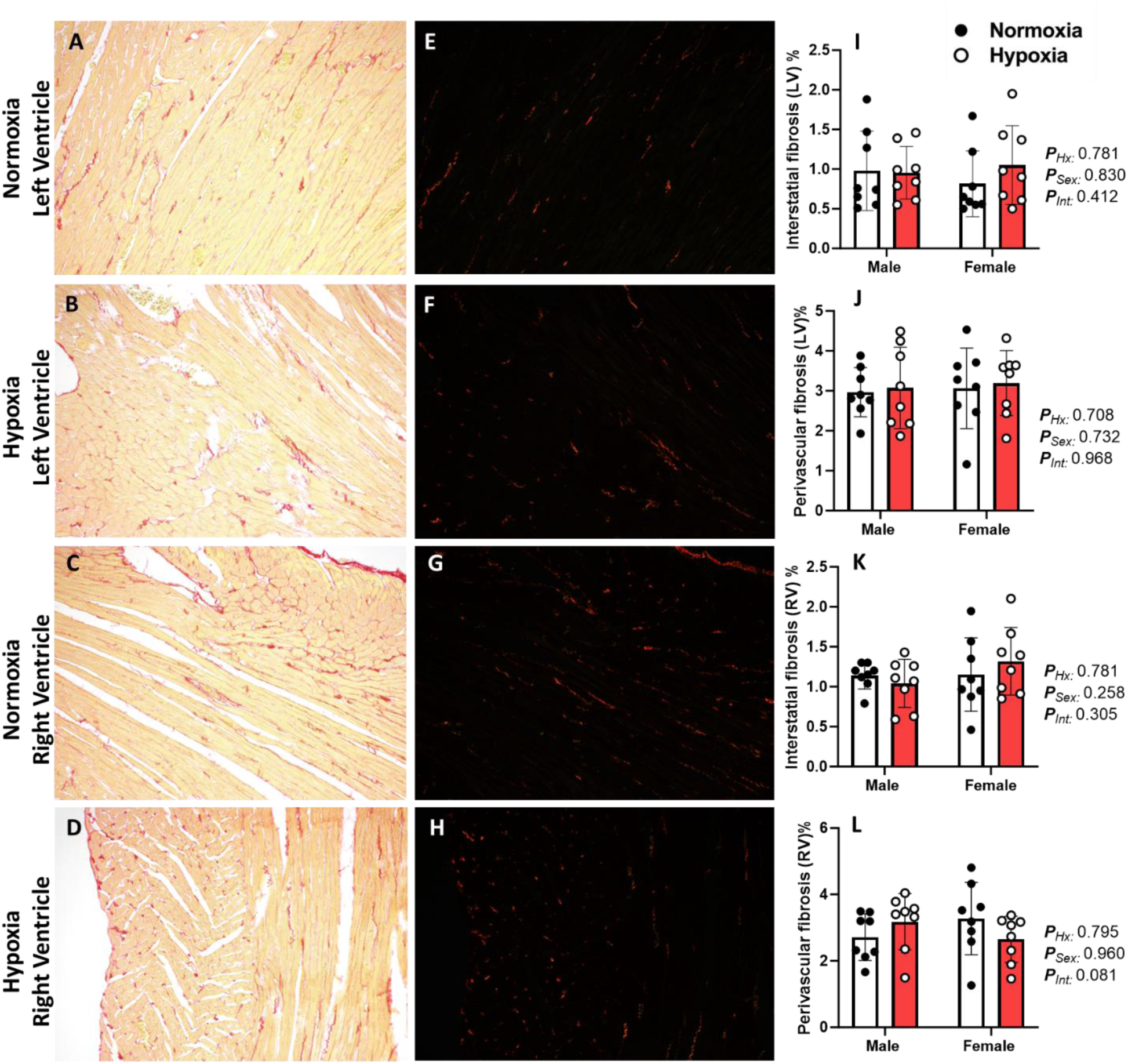
Histological analysis of cardiac fibrosis within the left and right ventricle. Picrosirius red staining revealed no significant difference in fibrosis between Normoxia and Hypoxia animals in either the left or right ventricle. Representative Brightfield (A, B, C, D) and Fluorescence (E, F, G, H) micrographs of cardiac tissue from offspring of normoxic and hypoxic pregnancies. Percentage interstitial (I, K) and perivascular fibrosis (J, L) from the left and right ventricle respectively. XX magnification, Scale = XX. Data analysed as a 2way ANOVA. Normoxia = closed circles, Hypoxia = open circles. *P*<0.05 was considered significant.

### Fetal hypoxia causes Ca^2+^ cycling abnormalities and AP prolongation

The increased arrhythmia susceptibility in offspring from hypoxic pregnancies was associated with abnormalities in the kinetics of the Ca^2+^ transient and the AP. In support of our previous findings^45^, male and female offspring from hypoxic pregnancies had prolonged Ca^2+^ transients in isolated hearts and isolated ventricular myocytes, as evidenced by an increased DT50 (Fig. 5A and Fig 6C), DT90 (Fig. 5B and Fig. 6E), and a prolonged decay time constant (tau) of the Ca^2+^ transient (Fig. 6G), compared to controls. Ca^2+^ transient rise time (Fig. 5C) was similar between groups, but the ventricular myocyte Ca^2+^ transient amplitude was reduced in adult offspring of hypoxic pregnancies (Fig. 6A, P=0.004), compared to controls. Male and female offspring from hypoxic pregnancies also had prolonged action potential duration in sinus rhythm, as evidenced by increased APD50 and APD90 in isolated hearts (P<0.05, Fig. 5E, F). There was no change in the amplitude of the caffeine transient, indicating no change in SR content (Fig. 6B), and no change in decay of the caffeine transient, other than an increase in DT50 in the fetal hypoxia group, compared to controls (Fig. 6D, P=0.035), indicating a potential reduction in NCX function. Lastly, the fetal hypoxia group significantly accumulated more Ca^2+^ during diastole when going from sinus rhythm to the start of arrhythmia, compared to controls (p=0.006), and this effect was worse in males than females (p=0.001, Fig. 5G).

**Figure 5:**
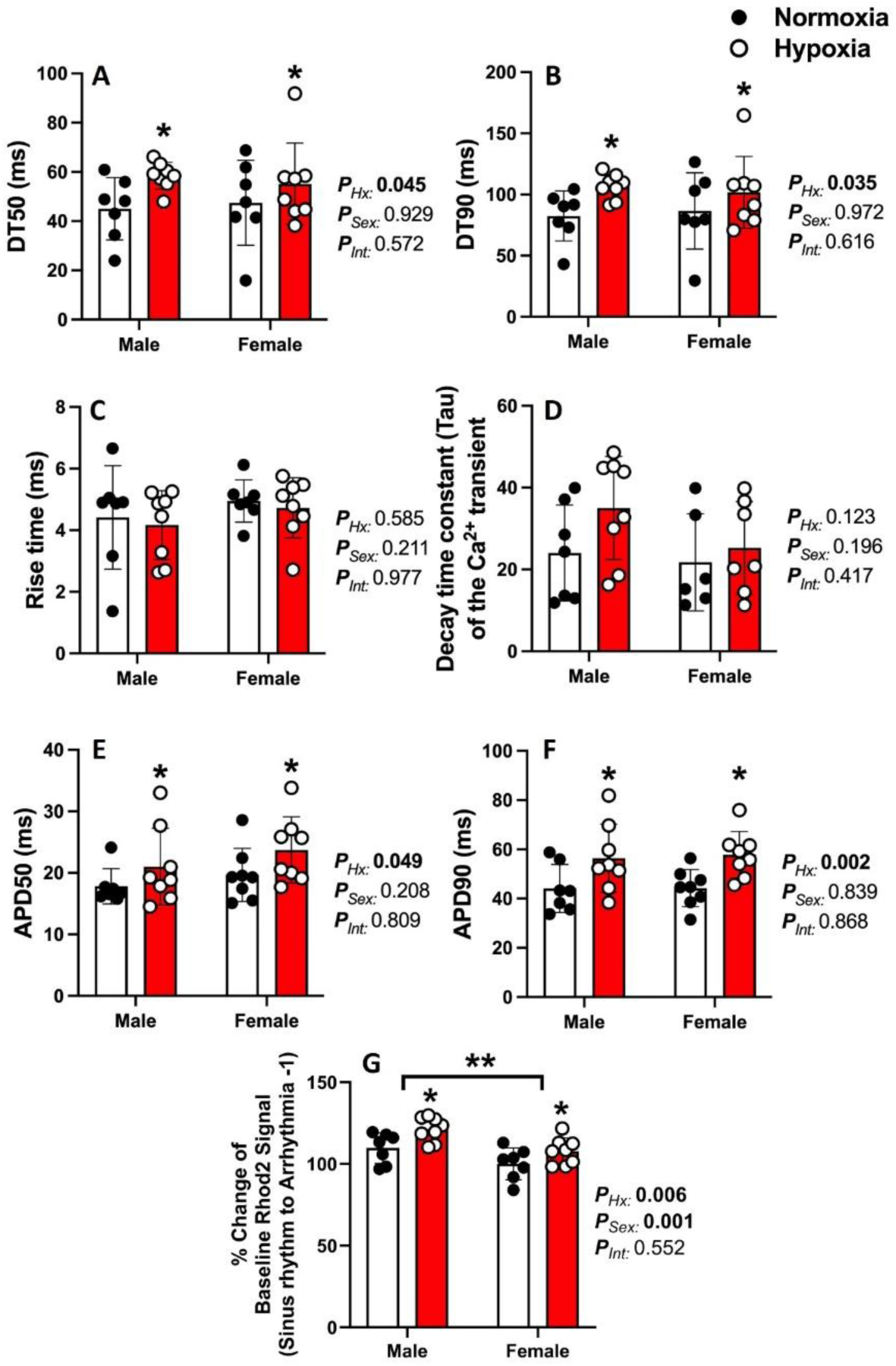
Calcium transient and action potential kinetics of isolated hearts under sinus rhythm. Data show the mean ± SD with each sample as dot plots for (A) Time taken for 50% reduction in Ca^2+^ from peak (DT50) and (B) 90% reduction in Ca^2+^ from peak (DT90). (C) Rise time of the calcium transient. (D) Time constant of the decay of the Ca^2+^ transient. (E) Action potential duration at 50%, and (F) 90%. (G) Percent change in baseline diastolic signal of Rhod2 between sinus rhythm and end stage of the arrhythmia protocol. Data analysed as a 2way ANOVA. Normoxia = closed circles, Hypoxia = open circles. *P*<0.05 was considered significant.

**Figure 6:**
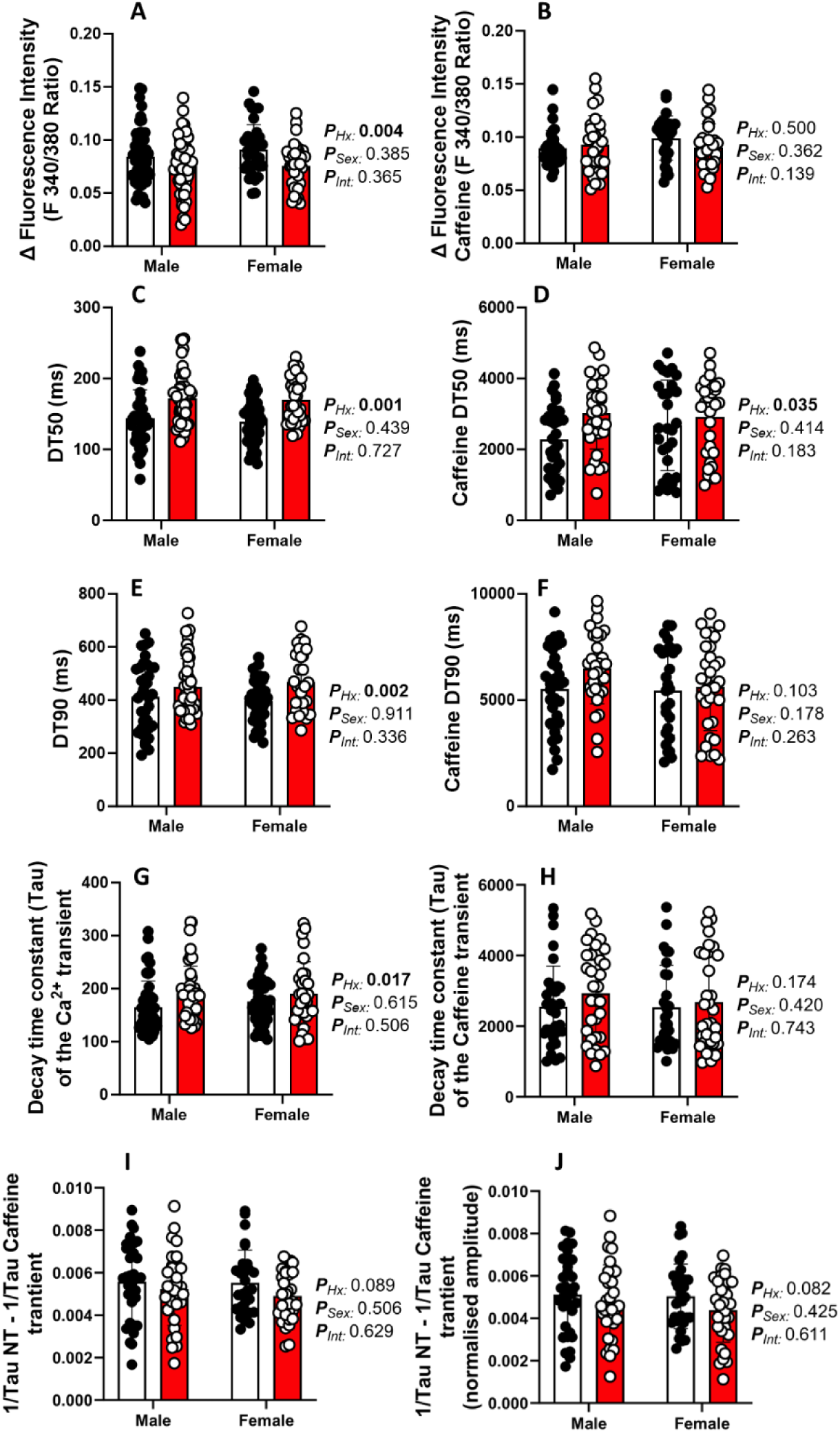
Systolic and diastolic Ca^2+^ and caffeine transient measures within isolated cardiomyocytes. A and B, systolic Ca^2+^ measure of peak fluorescence intensity in isolated cardiomyocytes from NT and caffeine transients, respectively. C and D, time for 50% decay of the Ca^2+^ and caffeine transients, respectively. E and F, time taken for isolated cells to relax to 50% of the amplitude of contraction from isolated cardiomyocytes from NT and caffeine transients, respectively. G and H, time constant for the decay of the Ca^2+^ transient in isolated cardiomyocytes from NT and caffeine transients, respectively. I and J are reciprocal decay of the calcium transient minus the decay of the caffeine transient, without and with normalisation to the amplitude of the calcium transient, respectively. Data analysed as a 2way ANOVA. Normoxia = closed circles, Hypoxia = open circles. *P*<0.05 was considered significant.

### Offspring from hypoxic pregnancies are more susceptible to ventricular myocyte calcium waves

We often observed spontaneous diastolic Ca^2+^ release events in the hypoxic group when hearts were burst paced, indicative of DADs (Fig. 2E). These events were also apparent in isolated ventricular myocytes, with offspring from hypoxic pregnancies having an increased percentage of ventricular myocytes that displayed Ca^2+^ waves (P<0.05, Fig. 7B) and Ca^2+^ sparks (Fig. 8) in the absence and presence of isoprenaline, compared to controls. Of the cells that underwent spontaneous Ca^2+^ release events, there were a higher number of these events per minute in cells isolated from offspring of hypoxic pregnancy compared to normoxic controls in both sexes (P<0.0001, Fig. 7C). Lastly, isolated myocytes from the fetal hypoxia group also had a relatively higher amplitude of Ca^2+^ waves, compared to normoxic controls (P=0.009, Fig. 7D), indicating more Ca^2+^ release from the SR during the wave.

**Figure 7:**
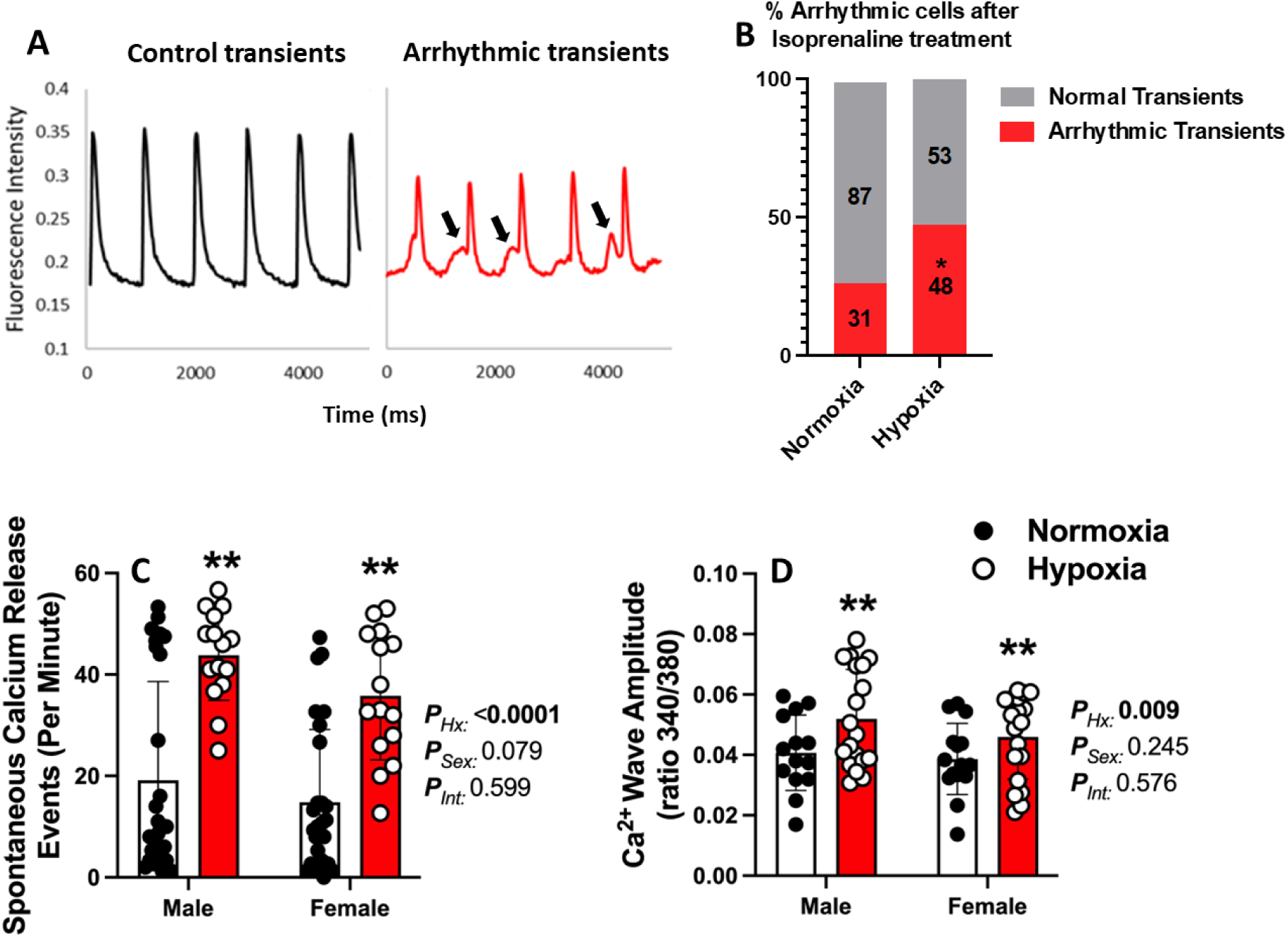
Spontaneous calcium release events in isolated cardiac myocytes from offspring of normoxic and hypoxic pregnancy. (A) Example Ca^2+^ transients from isolated cells from a cell behaving normally and one displaying spontaneous calcium release events after treatment with isoprenaline (arrows indicate calcium waves). (B) Percentage of cells demonstrating arrhythmic waves after isoprenaline (100nM) treatment in isolated myocytes from offspring of normoxic and hypoxic pregnancy. (C) Quantification of the number spontaneous calcium release events per minute in cells displaying calcium waves. (D) Amplitude of Ca^2+^ waves in cells that demonstrated spontaneous calcium release events. Data analysed as a 2way ANOVA. Normoxia = closed circles, Hypoxia = open circles. *P*<0.05 was considered significant.

**Figure 8.**
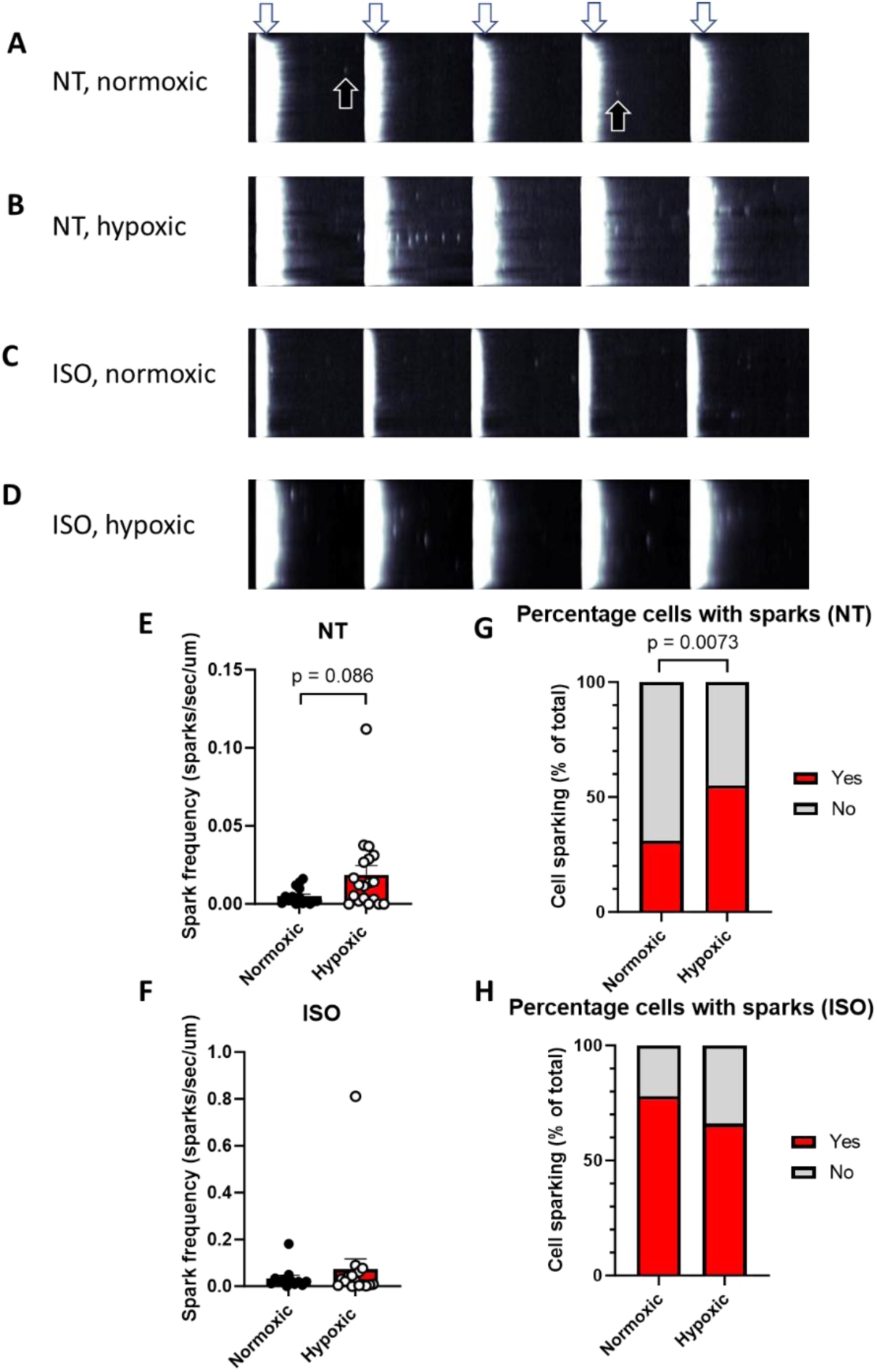
Fetal hypoxia increases the frequency of calcium sparks in isolated ventricular myocytes. Example linescan of myocyte from rats exposed to (A) Normoxia (NT), (B) Hypoxia (NT), (C) Normoxia (ISO) and (D) Hypoxia (ISO) during development. The white arrows demonstrate stimulation events and black arrows depict subsequent calcium sparks. Number of sparks measured per second per µm of cell length in myocytes from normoxic and hypoxic pregnancies subject to NT (E) and isoprenaline (100nM) (F). The percentage of cells where sparks were detected were normalised to the total number of cells under NT (G) and isoprenaline conditions (H).

### Expression of genes and proteins within the excitation-contraction coupling pathway

Fetal hypoxia resulted in increased mRNA expression of *RYR2* (P=0.0066, Fig. 9A) and decreased mRNA expression of *ATP2A2* (SERCA2a, P=0.0010, Fig. 9B) in the adult hearts of male offspring from hypoxic pregnancy. This change was also reflected in protein expression where SERCA2 expression (P=0.0144, Fig. 10I) and the ratio of SERCA2/PLN (P=0.0123, Fig. 10J) were both downregulated in the hearts of adult offspring from hypoxic pregnancy. There was no change in mRNA expression of any of the other measured genes in excitation-contraction coupling; *CAV1.2* (pore forming subunit of the L-type calcium channel), *PLN*, *CASQ* or *NCX* mRNA expression (Table 3, Fig. 9C, D, E, F). There was also no change in the cardiac protein expression of PLN, Phospho-PLN, cTnI, Phospho-cTnI, Calsequestrin-1, NCX1, LTCC, RYR2, Phospho-RYR2 and Connexin-43 between any experimental groups (Fig. 10B-H, K-P).

**Figure 9:**
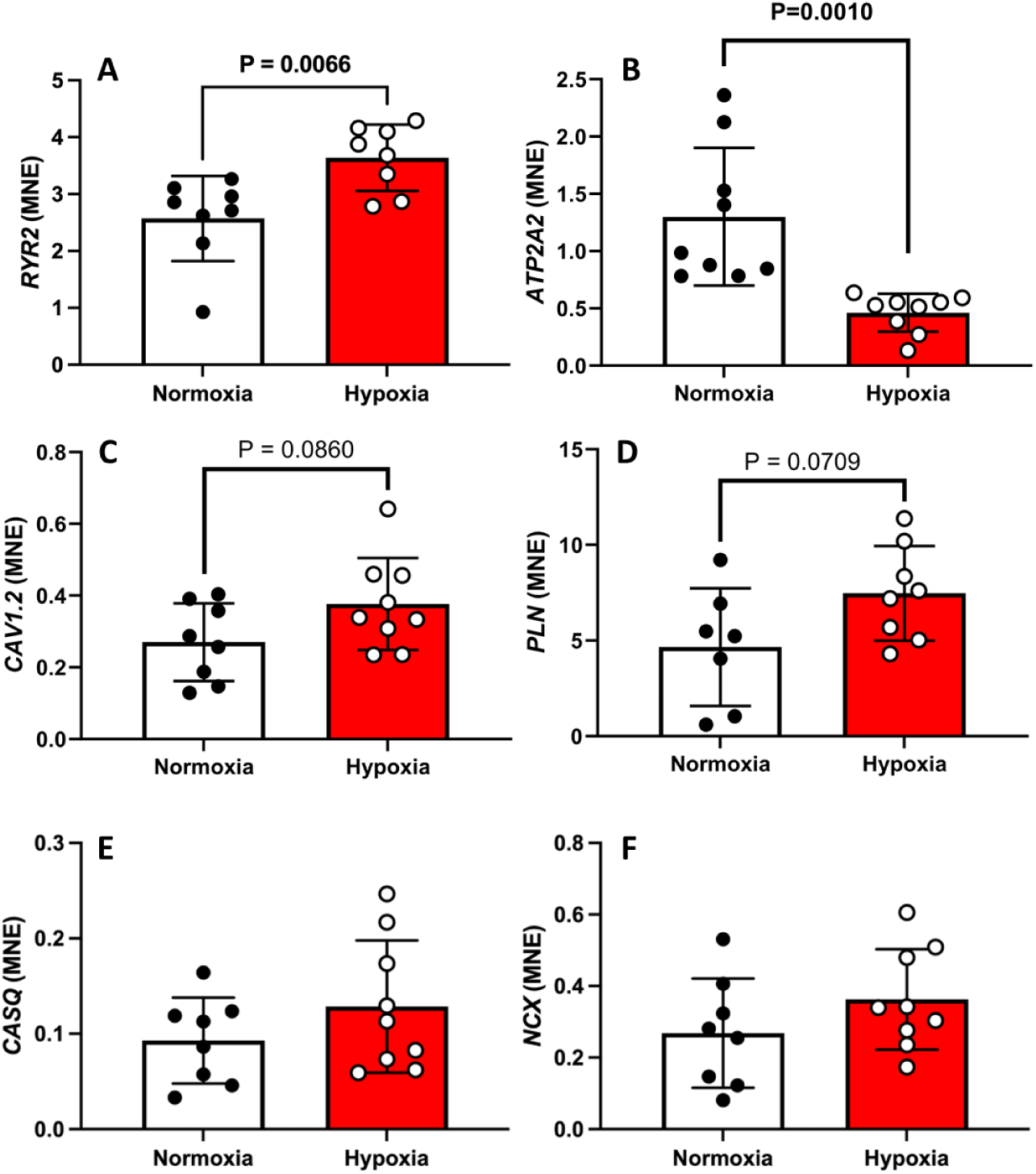
mRNA expression of core components of the excitation contraction pathway. Data show the mean ± SD with each sample as dot plots for the mRNA expression of *RYR2 (*A), *ATP2A2* (B)*, CAV1.2* (C), *PLN* (D), *CASQ* (E) and *NCX* (F). Data was analysed using an unpaired Student’s T-Test. P<0.05 was considered significant. Normoxia = closed circles, Hypoxia = open circles.

**Figure 10:**
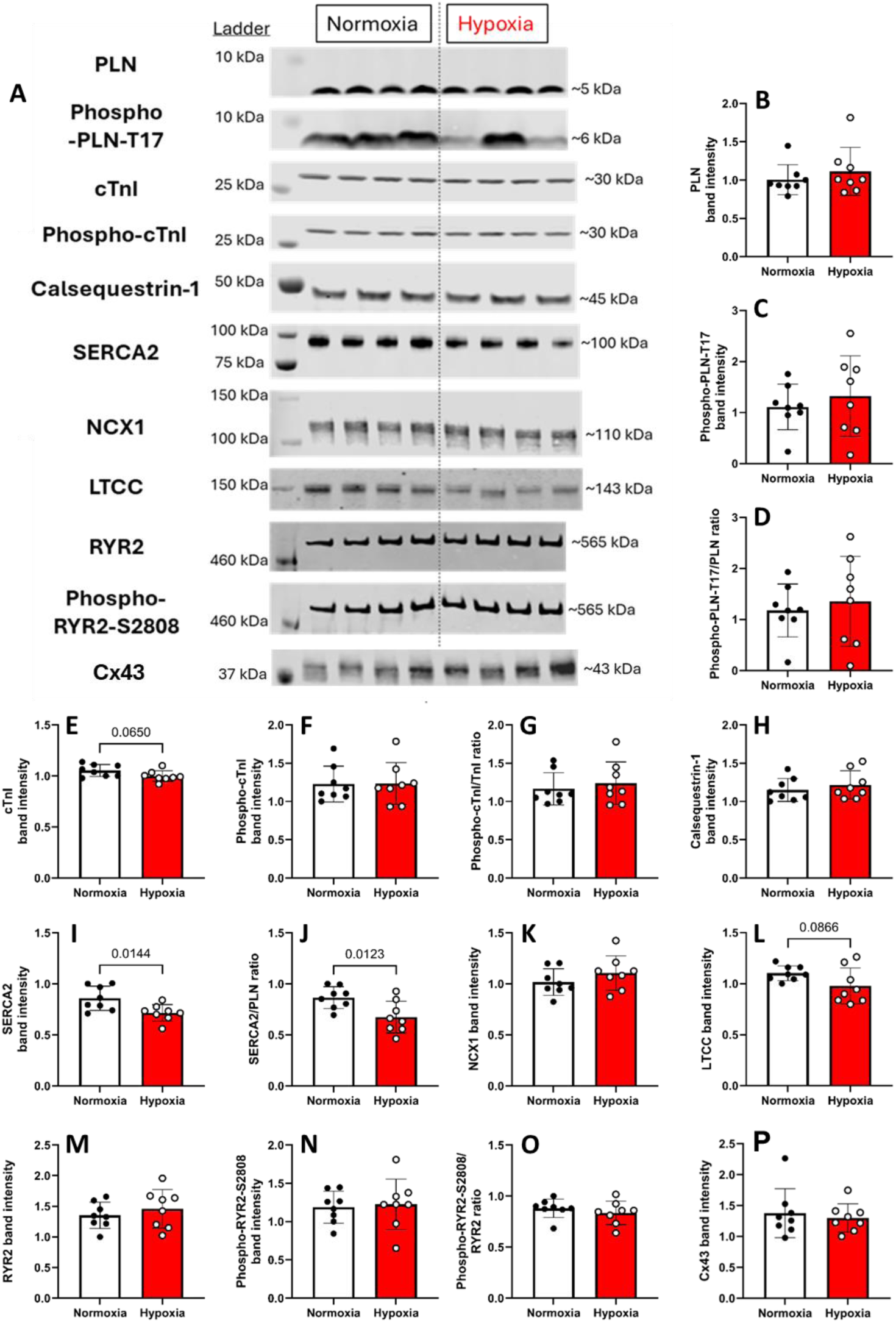
Protein abundance of core components of the excitation contraction pathway. Data show the mean ± SD with each sample as dot plots for the protein abundance of phospholamban (PLN, B), phospho-PLN-T17 (C), ratio of phospho-PLN-T17 to PLN (D), cardiac troponin I (cTnI, E), phospho-cTnI (F), ratio of phospho-cTnI to cTnI (G), calsequestrin-1 (H), SERCA2 (I), ratio of SERCA2 to PLN (J), sodium-calcium exchanger 1 (NCX1, K), L-type calcium channel (LTCC, L), ryanodine receptor 2 (RYR2, M), phospho-RYR2-S2808 (N), ratio of phospho-RYR2-S2808 to RYR2 (O) and Connexin43 (Cx43, P). Data was analysed using an unpaired Student’s T-Test for all proteins except PLN, cTnI, NCX and phospho-cTnI/cTnI ratio which used a Mann-Whitney test (data not normally distributed). P<0.05 was considered significant. Normoxia = closed circles, Hypoxia = open circles.

**Table 3:**
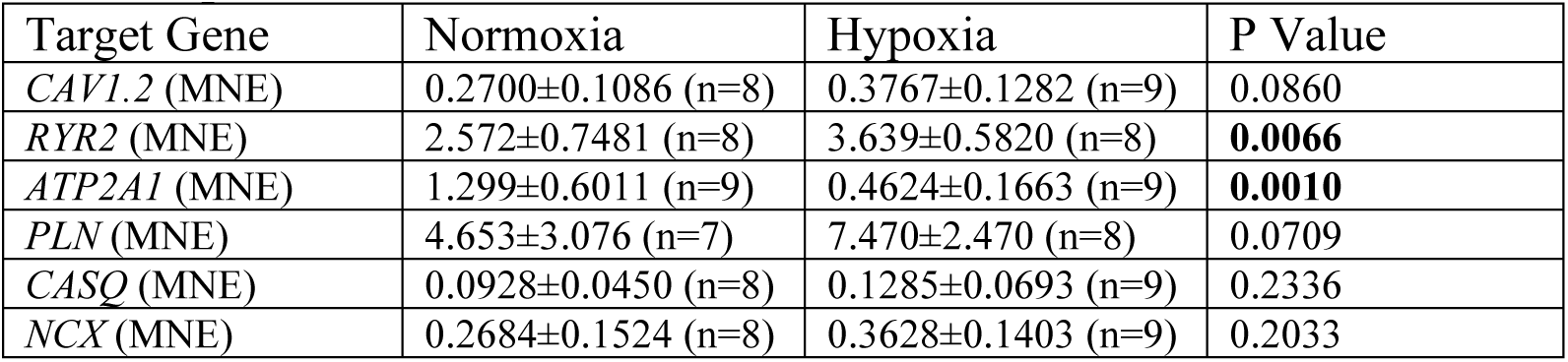
qRT-PCR measures.

## DISCUSSION

In this study, we show for the first time that ventricular arrhythmia sensitivity can be programmed by the intrauterine environment. Adult isolated hearts from rat offspring that experienced fetal hypoxia were significantly more likely to develop MVTs and PVTs under stress, and this was associated with abnormalities in excitation-contraction coupling that promote DADs and triggered activity.

### Fetal hypoxia increases offspring cardiac arrhythmia susceptibility

The hearts of offspring from hypoxic pregnancies were significantly more sensitive to arrhythmia, compared to controls. Indeed, most males from hypoxic pregnancies developed arrhythmia at low pacing frequencies in the absence of isoprenaline, and some even developed arrhythmias in sinus rhythm. Females from hypoxic pregnancies were also more susceptible to arrhythmia than controls, but slightly less sensitive than their male counterparts, with arrhythmias mostly developing with a combination of isoprenaline and burst pacing.

In both sexes, the arrhythmias generally manifested as PVT’s, resembling Torsade de Pointes (TdP), but some initially presented as MVT’s. MVT has often been associated with abnormal myocardial substrate, whereas PVTs are more common in acute and chronic ischemic heart disease. Interestingly some anti-arrhythmic drugs can ‘convert’ PVTs into MVTs, suggesting that the underlying substrate may also be important for PVTs. TdP often occurs in arrhythmia disease states such as Long QT Syndrome, which is associated with a prolonged AP duration, which we also observed in our model, indicating a potential common underlying mechanism.

### Mechanism driving arrhythmia susceptibility in offspring from hypoxic pregnancies

The presence of MVTs and PVTs suggest the presence of both reentry and triggered activity, respectively, in offspring from hypoxic pregnancies. Structural obstacles can often drive reentry circuits, and although previous studies have shown increased fibrosis in adult offspring from hypoxic pregnancies using 10.5% oxygen^46^, we did not find any change in collagen with our 13% oxygen model. Reentry arrhythmias can also be promoted by a reduction in electrical coupling, bought on by reduced expression of gap junction proteins (connexin 40 and 43^47^); but we found no change in conduction velocity or connexin expression between experimental groups. Alternatively, reentry circuits can occur via cellular heterogeneities in the AP refractory period^48^. In support of this concept, we found a prolongation of AP duration in the hearts of adult offspring from hypoxic pregnancies, which may increase the refractory period and predispose the ventricle to reentry circuits.

The presence of MVTs and PVTs in offspring from hypoxic pregnancies suggest the arrhythmias may have originated from both reentry and triggered activity, respectively. Re-entry depends on the relationship between CV and effective refractory period (ERP), such that the wavelength of the propagating impulse (CV × ERP) is short enough to permit the wavefront to re-excite adjacent recovered tissue. In the present study, we observed a strong trend towards a reduction in CV (p=0.052) in offspring from hypoxic pregnancies, which promotes re-entry by shortening the excitation wavelength, allowing a propagating wavefront to re-encounter recovered myocardium and sustain a functional re-entrant circuit. This was observed in the absence of fibrosis or gap-junction remodelling, which excludes structural discontinuity or impaired cell-to-cell coupling as primary drivers of arrhythmogenesis. However, reentry circuits can also occur via cellular heterogeneities in the AP refractory period^48^. Notably, offspring from hypoxic pregnancies also had a prolongation of the action potential, which can increase dispersion of repolarization and refractoriness, and predispose the heart to functional unidirectional block and stabilizing re-entrant circuits. These results suggest altered ionic currents, rather than architectural changes, could be responsible for an increased susceptibility to reentry arrhythmias in offspring previously exposed to fetal hypoxia.

In contrast to reentry, triggered arrhythmias are driven by early or delayed afterdepolarisations (EADs and DADs). Despite AP prolongation in hypoxic offspring, we did not find any evidence of EADs. However, we found an increase in the frequency of Ca^2+^ sparks, Ca^2+^ waves and DADs in isolated hearts and ventricular myocytes from individuals previously exposed to fetal hypoxia. DADs constitute a major arrhythmogenic mechanism in numerous cardiac diseases, including heart failure, catecholaminergic polymorphic ventricular tachycardia (CPVT) and right ventricular outflow tract tachycardia^35^. They are driven by spontaneous SR Ca^2+^ release during diastole that diffuse to contiguous junctional SR clusters to trigger propagating Ca^2+^ waves ^49^. These waves elevate intracellular calcium, activate the NCX, and give rise to a transient inward current which causes a depolarisation during diastole, leading to triggered activity. Therefore, our data suggests fetal hypoxia increases the likelihood of triggered activity by increasing the incidence of spontaneous Ca^2+^ release events and DADs.

### Mechanism driving spontaneous Ca^2+^ release and DADs

While the mechanism driving spontaneous Ca^2+^ release and Ca^2+^ waves in offspring from hypoxic pregnancies was not completely resolved in this study, our data suggest increased RyR leak, compromised Ca^2+^ clearance and reduced L-type Ca^2+^ channel expression may converge to promote DADs.

Spontaneous SR Ca^2+^ release (SR leak) occurs when SR Ca²⁺ content exceeds the threshold for RyR2 opening ^35,50^. Given that SR Ca^2+^ content was similar between experimental groups, it is likely that the increased SR leak in offspring from hypoxic pregnancies is driven by a reduced threshold for RyR opening (RyR sensitisation). This can occur under several circumstances, including post-translational modifications of RyR2 ^51^, a decrease in intraluminal SR Ca^2+^ buffering ^35^, and altered cytosolic Ca^2+^ buffering ^52^. Given that CASQ2 expression and phosphorylation was unchanged, our data would argue against SR luminal buffering as playing a role. From the perspective of posttranslational modifications, the phosphorylation status of RyR was unchanged, but our previous work on the same rodent model demonstrated that fetal hypoxia increases basal ROS production in the fetal and adult life stages ^24,25,53^. Increased mitochondrial ROS may therefore promote oxidative modification of RyR2, a mechanism known to reduce the threshold for channel opening and enhance diastolic SR Ca²⁺ leak.

In addition to an increased RyR leak, we found that offspring from hypoxic pregnancies had compromised Ca^2+^ clearance, manifested as prolonged Ca^2+^ transients bought about by reduced SERCA2A protein expression and activity. This phenotype was also observed in our previous study on the same rodent model^39^, and likely contributes to the diastolic dysfunction that characterises offspring from hypoxic pregnancies. In addition, a reduction in SERCA2A expression or activity decreases the buffering capacity of the SR, leading to higher local cytosolic Ca²⁺ following spontaneous SR Ca^2+^ release events. This facilitates propagation of Ca²⁺ waves by lowering the threshold for adjacent RyRs to be recruited via Ca²⁺-induced Ca²⁺ release. This effect is likely exacerbated by the reduction in troponin C protein expression that was also observed in offspring from hypoxic pregnancies, which further diminishes cytosolic Ca²⁺ buffering. Together, these changes create a cellular environment in which spontaneous Ca²⁺ release events are more likely to escalate into regenerative waves.

Despite the increased diastolic SR Ca²⁺ leak we observed, SR Ca²⁺ content was unchanged between experimental groups, implying that compensatory mechanisms must preserve SR loading. Interestingly, we found offspring from hypoxic pregnancies had a significantly smaller Ca^2+^ transient, with a strong trend towards reduced protein expression of the L-type Ca^2+^ channel. We have previously shown that a reduction in L-type Ca²⁺ current can paradoxically increase or maintain SR Ca²⁺ content under certain conditions^54^. This is because lowering the trigger for Ca²⁺–induced Ca²⁺ release reduces the fraction of SR Ca²⁺ released per beat and therefore diminishes Ca²⁺ efflux from the cell, allowing net Ca²⁺ balance to favour preservation of SR load^54^. Therefore, one interpretation of our data is that increased RyR2 sensitivity combined with reduced SERCA2A and troponin C-mediated buffering lowers the threshold for spontaneous SR Ca²⁺ release and increases the likelihood and propagation of Ca^2+^ waves. Reduced L-type Ca²⁺ channel expression helps to maintain SR Ca²⁺ content despite the leak, providing sufficient Ca²⁺ to propagate diastolic Ca²⁺ waves. These waves activate the NCX, generating delayed afterdepolarizations and promoting triggered arrhythmias.

This mechanism may be sufficient to trigger arrhythmias on its own, as evidenced by some males developing arrhythmias in sinus rhythm. However, as expected, arrhythmias in offspring from hypoxic pregnancies were more common at higher pacing frequencies and in the presence of isoprenaline. These conditions were also associated with elevated diastolic Ca²⁺ and prolonged action potentials. Therefore, SERCA2A activity may be sufficient to maintain normal diastolic Ca²⁺ levels under baseline conditions, but when pacing frequency is increased or in the presence of isoprenaline (which increases SR Ca^2+^ load), the time available for Ca²⁺ reuptake is reduced, compromising cytosolic buffering. Under these conditions, elevated SR Ca²⁺ load, RyR2 sensitisation, and diminished cytosolic buffering synergize to facilitate the initiation and propagation of diastolic Ca²⁺ waves.

### Sex-dependent differences

In adulthood, pre-menopause females are generally considered to be more protected from cardiovascular disease than males, possibly due to the presence of estrogen ^55,56^. The higher levels of androgens in males (such as testosterone) may also contribute to sex-dependent differences, as they are thought to be pro-hypertensive. Previous studies have also shown that females appear to be less sensitive than males to cardiac programming. These dimorphic responses to manipulations during embryonic and fetal development may arise from differences in the ability of each sex to respond and adapt to specific challenges ^57,58^. Data from this study supports this contention, because hearts from female rats from hypoxic pregnancy were significantly less likely to experience arrhythmia during burst pacing and isoprenaline treatment, compared to males. Furthermore, the kinetics of the Ca^2+^ transient and AP in isolated hearts were less affected by fetal Hypoxia in females versus males. Nevertheless, within our isolated cell experiments there were no differences observed between sexes. It is therefore likely that cardiac structural changes and/or connectivity between cells (for example expression of connexins) may play a large role in the sexual dimorphic response we observed, which were not present within isolated cells.

### Conclusions and Future Directions

This study provides the first direct evidence that susceptibility to ventricular arrhythmia can be programmed during fetal development; revealing a previously unrecognised developmental origin for arrhythmia. This has important clinical implications, as individuals exposed to hypoxic pregnancy complications (e.g. preeclampsia, placental dysfunction, anaemia, or high-altitude gestation) may carry a latent arrhythmic risk that is not captured by current risk stratification approaches. Critically, these data highlight pregnancy as a key window for primary prevention of adult arrhythmic disease. Given the central role of oxidative stress in developmental hypoxia–induced cardiac programming, strategies aimed at reducing fetal oxidative stress represent a promising avenue for early-life intervention to lower lifelong arrhythmia risk.

While the mechanism for increased arrhythmia sensitivity is still being resolved, it is clear that fetal hypoxia programmes abnormalities in excitation-contraction coupling that promote spontaneous Ca^2+^ release, Ca^2+^ waves, DADs and triggered activity. Our data also suggest the mechanism driving this phenotype involves increased RyR leak, compromised Ca^2+^ clearance (reduced SERCA expression) and reduced L-type Ca^2+^ channel expression. In order to definitively establish the source of DAD’s in offspring from hypoxic pregnancies, future work should assess RyR2 redox state and post-translational modifications, quantify the impact of mitochondrial-targeted antioxidants on SR Ca²⁺ leak, measure L-type Ca^2+^ channel density and kinetics, and evaluate changes in luminal and cytosolic Ca²⁺ buffering proteins. It would also be interesting to investigate epigenetic signatures, as changes in protein expression of EC coupling proteins could be due to epigenetic programming of genes during fetal life ^59^. Indeed, previous work has shown fetal hypoxia not only induced DNA methylomic and transcriptomic changes in the fetal hearts, but also had a delayed and lasting effect on the adult offspring hearts. This may help to explain why our data which shows fetal hypoxia increases protein and mRNA expression of *SERCA2a*.

## ABBREVIATION LIST

CICR: Ca^2+^-induced Ca^2+^ release
CVD: Cardiovascular disease
ECC: Excitation-contraction coupling
GD: gestational day
LV: left ventricle
NCX: sarcolemmal Na+/Ca2+ exchanger
NO: Nitric oxide
PKA: protein kinase A
RV: right ventricle
SERCA2A: sarcoplasmic reticulum Ca^2+^-ATPase
SI: sphericity index
MVT: monomorphic ventricular tachycardia
PVT: polymorphic ventricular tachycardia
VF: ventricular fibrillation
RyR: Ryanodine receptor.

## Acknowledgments

This work was funded by two British Heart Foundation (BHF) project grants awarded to GG, AWT and DG; Grant Numbers: PG/18/5/33527 & PG/23/11296.

## SUPPLEMENTARY FIGURES

**Figure S1:**
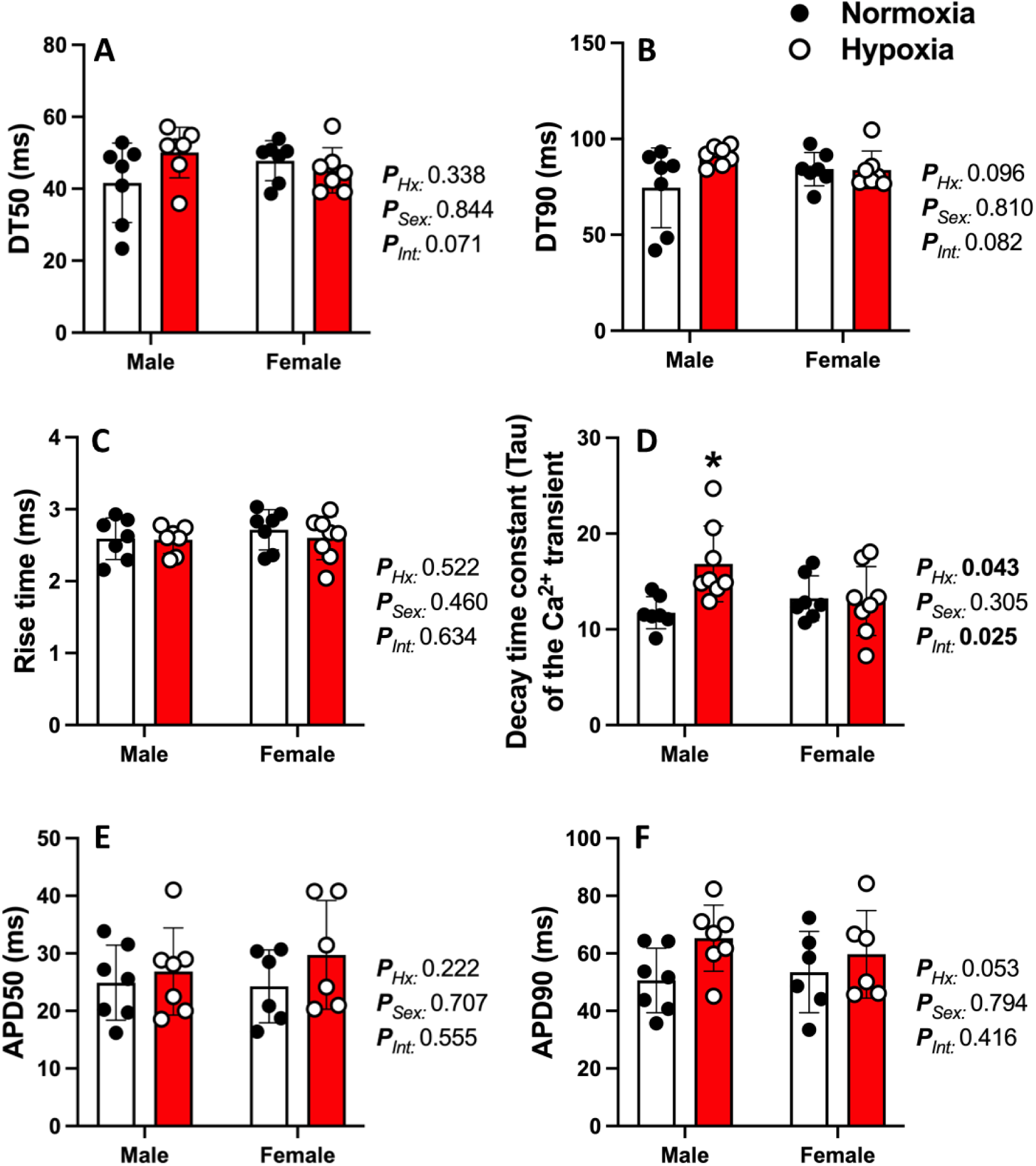
Calcium transient and action potential kinetics of isolated hearts under 5Hz stimulation. Data show the mean ± SD with each sample as dot plots for (A) Time taken for 50% reduction in Ca^2+^ from peak (DT50) and (B) 90% reduction in Ca^2+^ from peak (DT90). (C) Rise time of the calcium transient. (D) Time constant of the decay of the Ca^2+^ transient. (E) Action potential duration at 50%, and (F) 90%. Data analysed as a 2way ANOVA. Normoxia = closed circles, Hypoxia = open circles. *P*<0.05 was considered significant.

**Figure S2:**
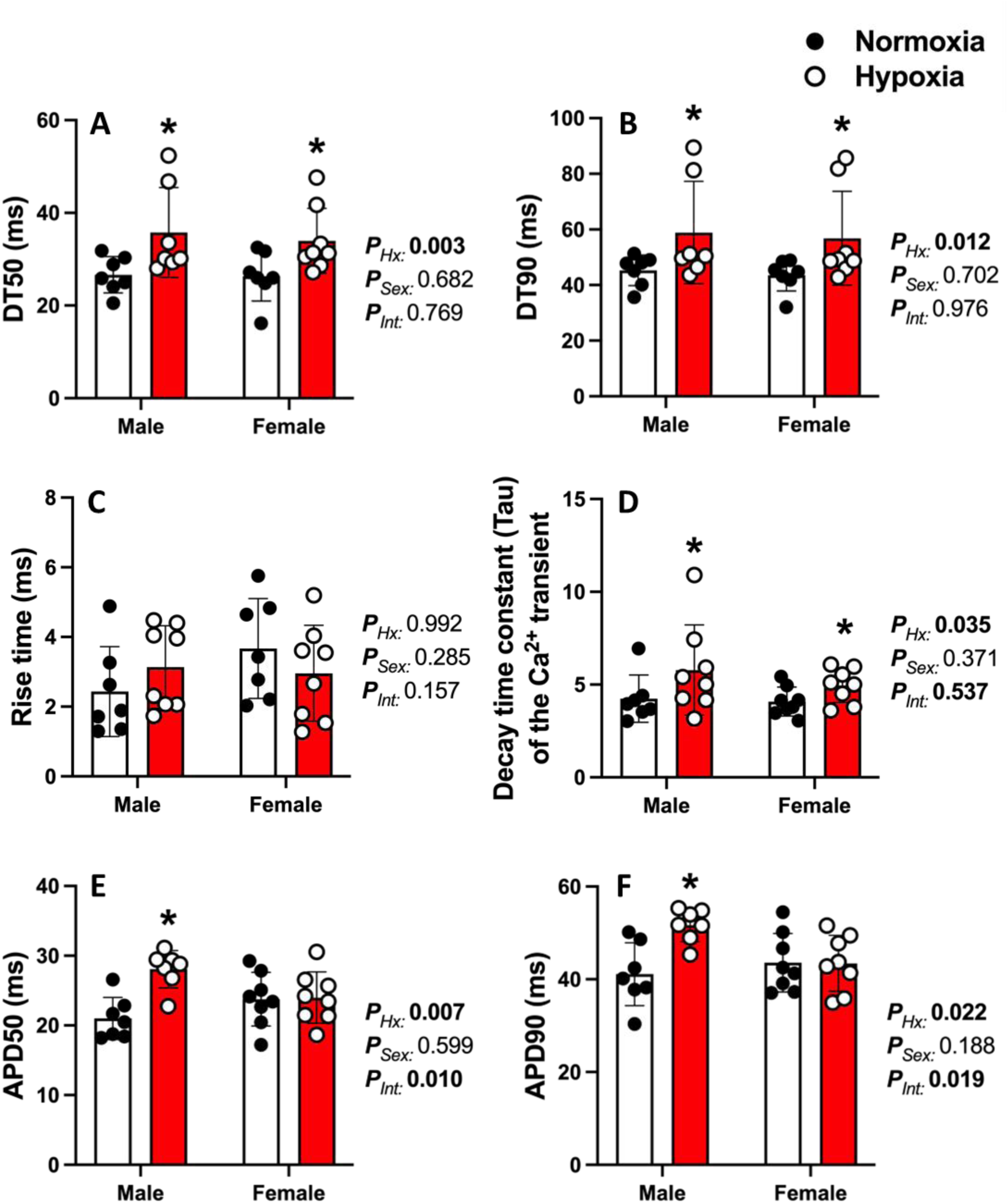
Calcium transient and action potential kinetics of isolated hearts under 10Hz burst pacing. Data show the mean ± SD with each sample as dot plots for (A) Time taken for 50% reduction in Ca^2+^ from peak (DT50) and (B) 90% reduction in Ca^2+^ from peak (DT90). (C) Rise time of the calcium transient. (D) Time constant of the decay of the Ca^2+^ transient. (E) Action potential duration at 50%, and (F) 90%. Data analysed as a 2way ANOVA. Normoxia = closed circles, Hypoxia = open circles. *P*<0.05 was considered significant.

